# Discovery of T-dioxygenases in bacteriophages and identification of a subclass that is dependent on a regulator protein for oxidation

**DOI:** 10.1101/2025.08.14.668559

**Authors:** Katherine H. O’Toole, Auriane Bouchet, Mia L. DeSanctis, Sean R. Lund, Sabaa Belkadi, Harold W. Bell, Lana Saleh

## Abstract

5-Methylpyrimidine dioxygenases (5mYOXs) are iron (II)/2-oxoglutarate-dependent enzymes that catalyze the oxidation of DNA 5-methylpyrimidines. Key members include mammalian ten-eleven translocation (TET) dioxygenases and J-base binding proteins (JBP) from trypanosomes, which oxidize 5-methylcytosine (5mC) and thymine (T) on DNA, respectively, and are essential in gene regulation. Using sequence similarity networks and genome mining, we highlight functional predictions within the 5mYOX superfamily and identify thousands of bacteriophage-derived sequences, often found in operons encoding DNA modification machinery and binding proteins. Through high-throughput in vivo functional assays, we confirm that T oxidation occurs in bacterial viruses, establishing the presence of T dioxygenases outside of eukaryotes. We provide the first evidence for a 5mYOX subclass that is inactive unless co-expressed with a regulator protein, and through structural modeling, show that this regulator bears homology to the bacterial partition protein B (ParB). We propose a model of interaction between ParB and 5mYOX that includes complex formation, with 5mYOX binding DNA and ParB utilizing CTP, as in bacteria, to form an optimal structural configuration for productive oxidation and possibly migration on the DNA. We show these enzymes retain key catalytic residues found in TET/JBP enzymes and employ AlphaFold2-guided mutagenesis to identify clade-specific features critical for T oxidation, including a variable insertion important for ParB-independent activity and a conserved C-terminal extension essential for T oxidation in ParB-dependent homologs. These findings uncover modular determinants, regulatory mechanisms, and the evolutionary diversity of T-dioxygenases, expanding the functional landscape of the 5mYOX superfamily.

**Significance:** This study demonstrates that enzymatic oxidation of thymidine (T) by 5-methylpyrimidine dioxygenases (5mYOXs) occurs in nature outside eukaryotes. Through bioinformatics and *in vivo* screening, we elucidate minimal T-dioxygenases that share conserved catalytic properties with eukaryotic 5mYOXs, yet possess unique domains, lineage-specific inserts, and/or regulators essential for activity. We uncovered two distinct, bacteriophage-encoded T-dioxygenase subclasses: one requiring a partition protein B (ParB)-like regulator for oxidation and another independent of it. This regulator is predicted to structurally resemble bacterial segregation ParB, which utilizes CTP binding and hydrolysis to migrate along DNA, suggesting novel mechanisms for T oxidation in phages. Our identification of subclass-specific features provides a foundation for further investigation of substrate selectivity, oxidation regulation, and 5mYOX divergence across domains of life.

---

Methylation and hydroxymethylation of pyrimidines at the C5 position are pivotal for orchestrating epigenetic regulation and safeguarding genomic integrity (1–4). These modifications are introduced either pre-replication via thymidylate synthase homologs or post-replication at the polymer level through the enzymatic actions of S-adenosylmethionine (SAM)-dependent DNA-(cytosine C5)-methyltransferases (C5-MTs) and 5-methylpyrimidine dioxygenases (5mYOXs) (3, 5–7). The 5mYOX superfamily, distinguished by its conserved jelly-roll fold, facilitates the oxidation of 5-methylcytosine (5mC) or thymine (T) on DNA in an iron (II) and 2-oxoglutarate (2OG)-dependent manner (8–12), encompassing both mammalian ten-eleven translocation (TET) dioxygenases and trypanosome J-base binding proteins (JBPs). Mammalian TET enzymes sequentially oxidize 5mC to 5-hydroxymethylcytosine (5hmC), 5-formylcytosine (5fC), and 5-carboxycytosine (5caC) (Fig. 1A), ultimately enabling DNA demethylation through base excision repair, thereby resetting the epigenetic mark to cytosine (13–15). These processes are integral to epigenetic regulation and cell fate determination (16–21). Similarly, trypanosome JBPs oxidize T to 5-hydroxymethyluracil (5hmU), which undergoes further modification to form Base J, 5-(β-D-glucosyloxymethyl)uracil, a key player in gene silencing mechanisms (Fig. 1B) (1, 22).

**Fig. 1.**
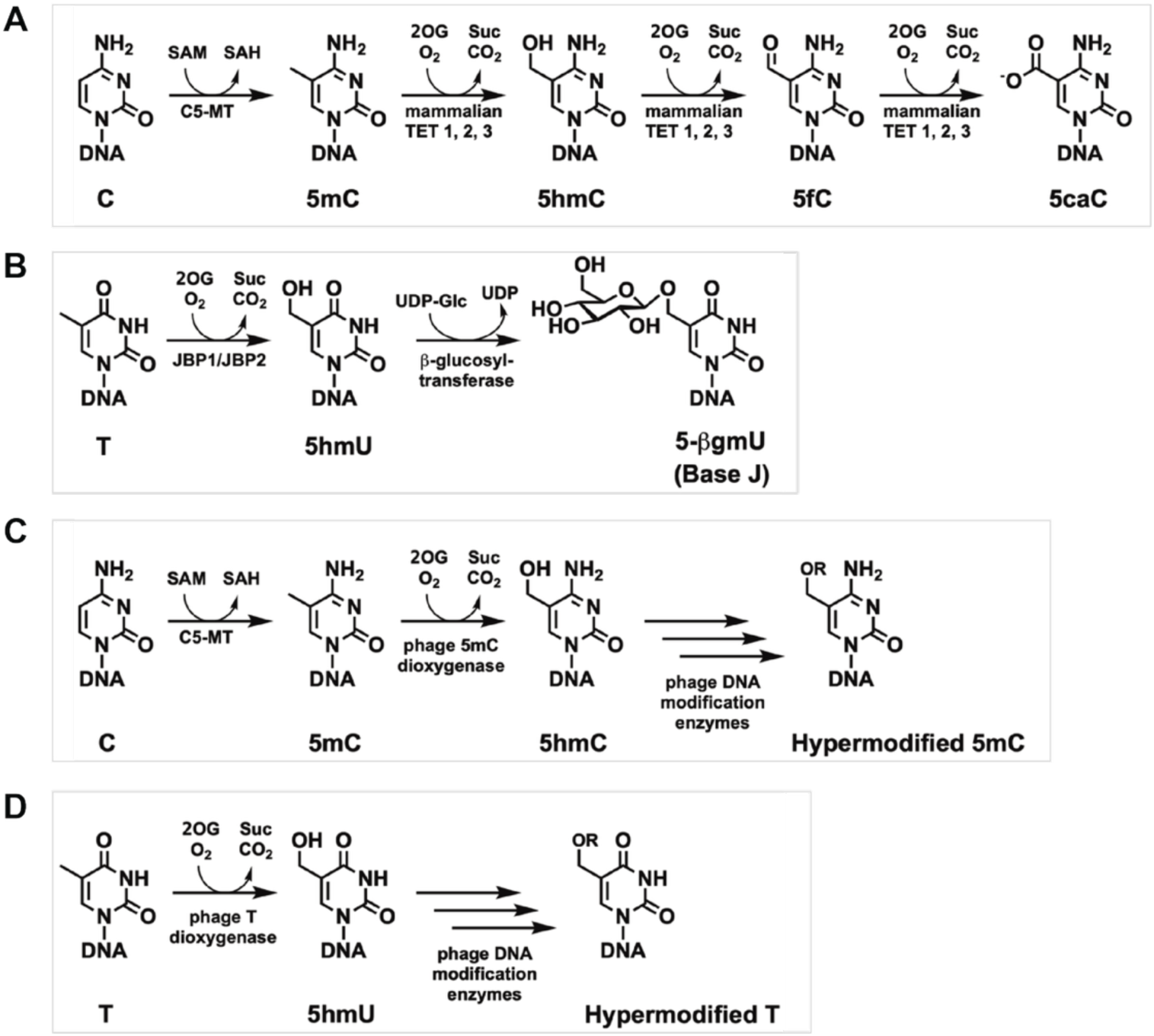
Reaction schemes for 5mYOX oxidation and DNA hypermodification. *(A)* Methylation of cytosine by SAM-dependent C5-MT and the iterative oxidation of 5mC by the mammalian TET dioxygenases 1-3 in epigenetic regulation of gene expression; Biosynthetic pathways for *(B)* Base J incorporation into DNA by JBP1/JBP2 and β-glucosyltransferase in parasitic trypanosomes; *(C)* formation of hypermodified C in phage by C5-MT, 5mC-dioxygenases, and downstream hypermodification enzymes; and *(D)* formation of hypermodified T in phage by T-dioxygenases and downstream hypermodification enzymes. S-adenosyl-homocysteine = SAH; Succinate = Suc.

Our group previously identified phage-encoded 5mYOXs capable of oxidizing 5mC to 5hmC, unveiling novel post-replicative DNA hypermodification pathways that yield a variety of sugar-modified cytosine derivatives (Fig. 1C) (7, 23). Notably, one enzyme, 5mYOX89, demonstrated a distinct preference for T over 5mC, marking it as the first post-replicative T-dioxygenase identified apart from JBP (7). Conversely, certain 5mYOXs, including 5mYOX67 and 5mYOX79, showed no oxidation activity towards 5mC or T. Subsequent analysis revealed these enzymes are associated with a gene encoding a partition protein B (ParB)-like protein, a member of the ParB superfamily (PF02195) (24, 25), which typically functions in bacterial chromosome partitioning through its nucleotide-binding domain (26, 27). Although the potential role of ParB in DNA modification pathways had been suggested (28, 29), it remained experimentally unconfirmed.

In this study, we employed genome mining and sequence similarity network (SSN) (30, 31) approaches to identify 5mYOX-containing biosynthetic gene clusters (BGCs) devoid of C5-MT genes, postulating that these enzymes might target T. We experimentally validated multiple phage-origin T-dioxygenases, categorizing them into two subclasses: regulator-dependent and - independent. Through a “mix and match” strategy, we assessed the specificity of ParB homologs to their respective 5mYOXs and explored the promiscuity observed in certain ParBs. Additionally, using multiple sequence alignment (MSA) and structural bioinformatics, we predicted domains unique to each T-dioxygenase subclass and probed their functional significance through deletion studies and protein chimera approaches. We find that 5mYOX-associated ParBs are divergent members of the ParB family lacking the DNA-binding domain but retaining features of the N-terminal nucleotide-binding domain and the C-terminal dimerization domain. Similarly, phage 5mYOXs share conserved catalytic features with their eukaryotic counterparts but exhibit unique substitutions and compact architecture that likely affect their substrate selectivity.

This study significantly advances our understanding of the biologically important 5mYOX superfamily and provides the first detailed characterization of phage T-dioxygenases, linking them to DNA hypermodification pathways (Fig. 1D). It demonstrates domains essential for T oxidation in phages and reports the first-ever regulator-dependent 5mYOX, drawing parallels to bacterial ParB’s CTP-dependent switch that governs DNA loading and migration (32, 33).

## Results

### Curation and SSN analysis of 5mYOX superfamily for T-dioxygenase candidate selection

We performed genome mining using the hidden Markov model (HMM) profile for the Pfam TET/JBP superfamily (PF12851) against the IMG/VR3 metaviriome database (34–36), identifying 2,323 non-redundant 5mYOX sequences. We then combined these with sequences annotated within the InterPro family (IPR024779) and manually curated them to construct SSNs that describe the 5mYOX superfamily using the Enzyme Function Initiative-Enzyme Similarity (EFI-EST) tool (30, 31) (Figs. 2 and 3). All-by-all pairwise alignments were conducted using *E*-value thresholds of 1 × 10^-35^, 1 × 10^-50^, and 1 × 10^-70^. The 40 % representative node network (*E*-value: 1 × 10^-35^) included 6,381 non-redundant sequences (< 100 % identity), comprising 3,111 nodes and 1.7 million edges, which coalesced into 6 independent clusters and 23 singleton clusters (Fig. 2). The superfamily’s distribution across domains of life was 63 % viral, 36 % eukaryotic, 1.5 % bacterial, and < 1 % archaeal (Fig. 2A). The most prominent cluster consisted exclusively of eukaryotic members within the Opisthokonta kingdom, predicted to be TET homologs (5mC-dioxygenases). As a first step towards selecting and functionally screening novel 5mYOXs, we examined the clustering of previously characterized enzymes, such as mammalian TETs, including mouse TET2 and human TETs 1-3 (5mC-dioxygenase, cyan squares), which cluster separately from phage-encoded 5mC-dioxygenases (green squares) (Fig. 2B), consistent with their functional differences as phage enzymes like 5mYOX43 are GpC-specific and stall at 5hmC (7). We next identified that “5mYOX89” (phage T-dioxygenase, purple square) and trypanosome JBPs (eukaryotic T-dioxygenases, orange squares) cluster together, with trypanosome JBPs forming a smaller sub-cluster (Fig. 2B). The larger cluster contains numerous sequences that we here predict to be T-dioxygenases (purple and magenta squares) based on the lack of activity toward 5mC (by a few characterized members (yellow squares)) (7), absence of a proximal C5-MT gene (Fig. 3), and their clustering close to trypanosome JBPs (10, 22) and 5mYOX89 (7) (Fig. 2B; *E*-value: 1 × 10^-35^, dotted black circle). Notably, the trypanosome JBPs connect to this larger cluster by only one edge, emphasizing the distant relation to predicted bacteriophage T-dioxygenases. Increasing the *E*-value threshold to 1 × 10^-50^ further resolved the clustering, partitioning JBPs (orange squares) and 5mYOX89 homologs (purple squares) into distinct sub-clusters. At *E*-value: 1 × 10^-70^, 5mYOX67 and 79, previously shown to be inactive towards 5mC or T (7) diverged into singleton nodes, indicating their low identity with other sequences.

**Fig. 2.**
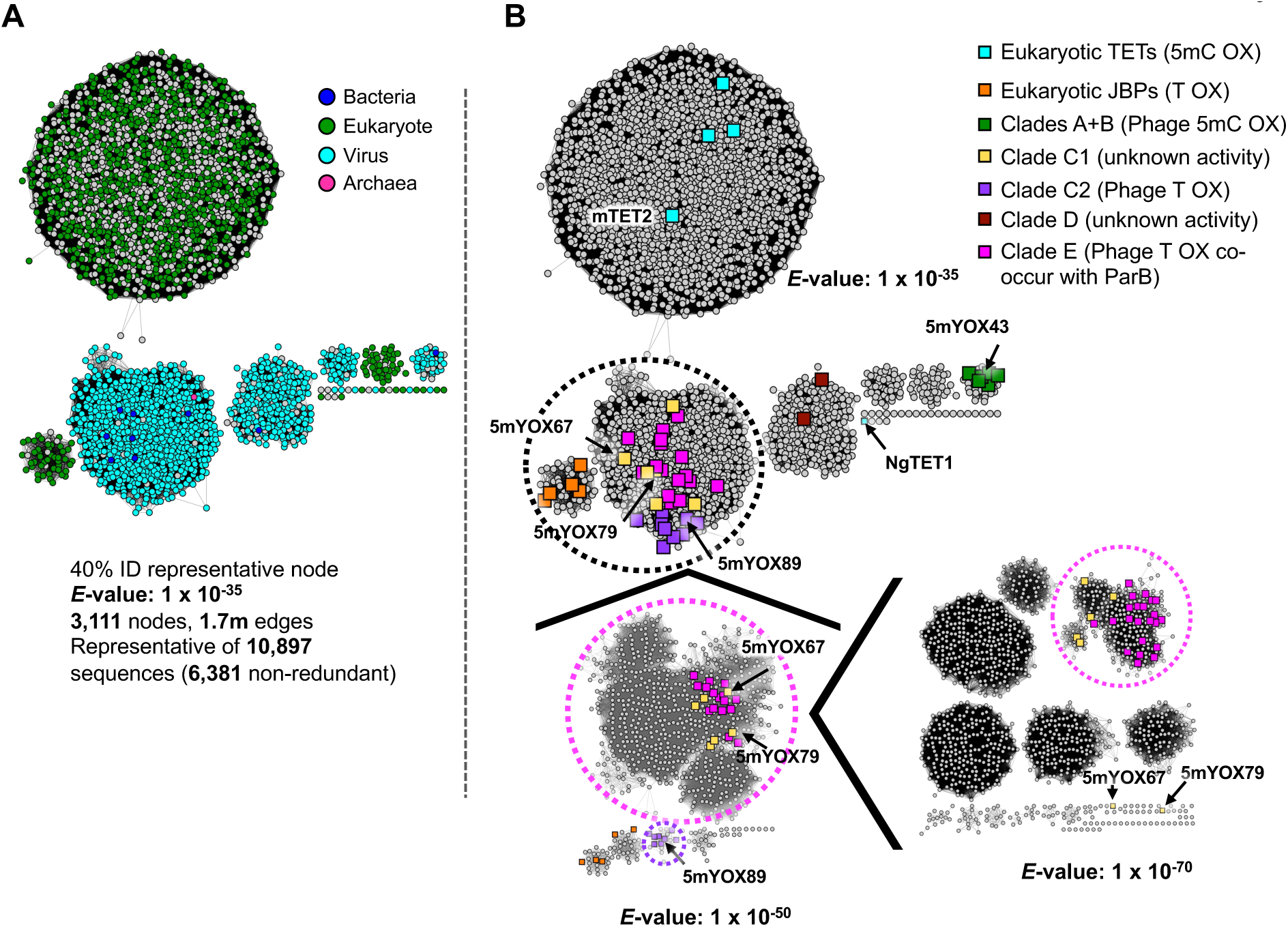
SSN representation of 5mYOX superfamily and selection of priority candidates for screening. *(A)* SSN 40 % representative node network of 5mYOX superfamily (*E*-value: 1 × 10^-35^) from 6,381 non-redundant sequences colored by domain of life: bacteria (blue), eukaryote (green), virus (cyan), and archaea (magenta). *(B)* SSNs colored by function and clade assignment: eukaryotic TETs (cyan), JBPs (orange), phage 5mC-dioxygenases Clades A/B (7) (green), Clade C1 (7) (yellow), Clade C2 phage T-dioxygenase (7) (purple), Clade D (7) (brown), and Clade E co-occurring with ParB (magenta). Characterized eukaryotic TETs highlighted: human hTET1-3, mouse mTET2, *Naegleria gruberi* NgTET1 (UniProt IDs: Q8NFU7, Q6N021, O43151, Q4JK59, D2W6T1, respectively). Characterized eukaryotic JBPs highlighted: *Leishmania major* LmJBP 1 and 2; *Trypanosoma brucei* TbJBP 1 and 2, and *Leishmania tarentolae Lt*JBP1 and 2 (UniProt ID: Q4QHM7, Q4QFY1, P86938, Q57X81, Q9U6M1, B6EU02, respectively). Purple dotted circle (*E*-value: 1 × 10⁻^50^; 40 % representative node) and magenta dotted circle (*E*-value: 1 × 10⁻^70^; 70 % representative node) represent sequences that diverged away from trypanosome JBPs.

**Fig. 3.**
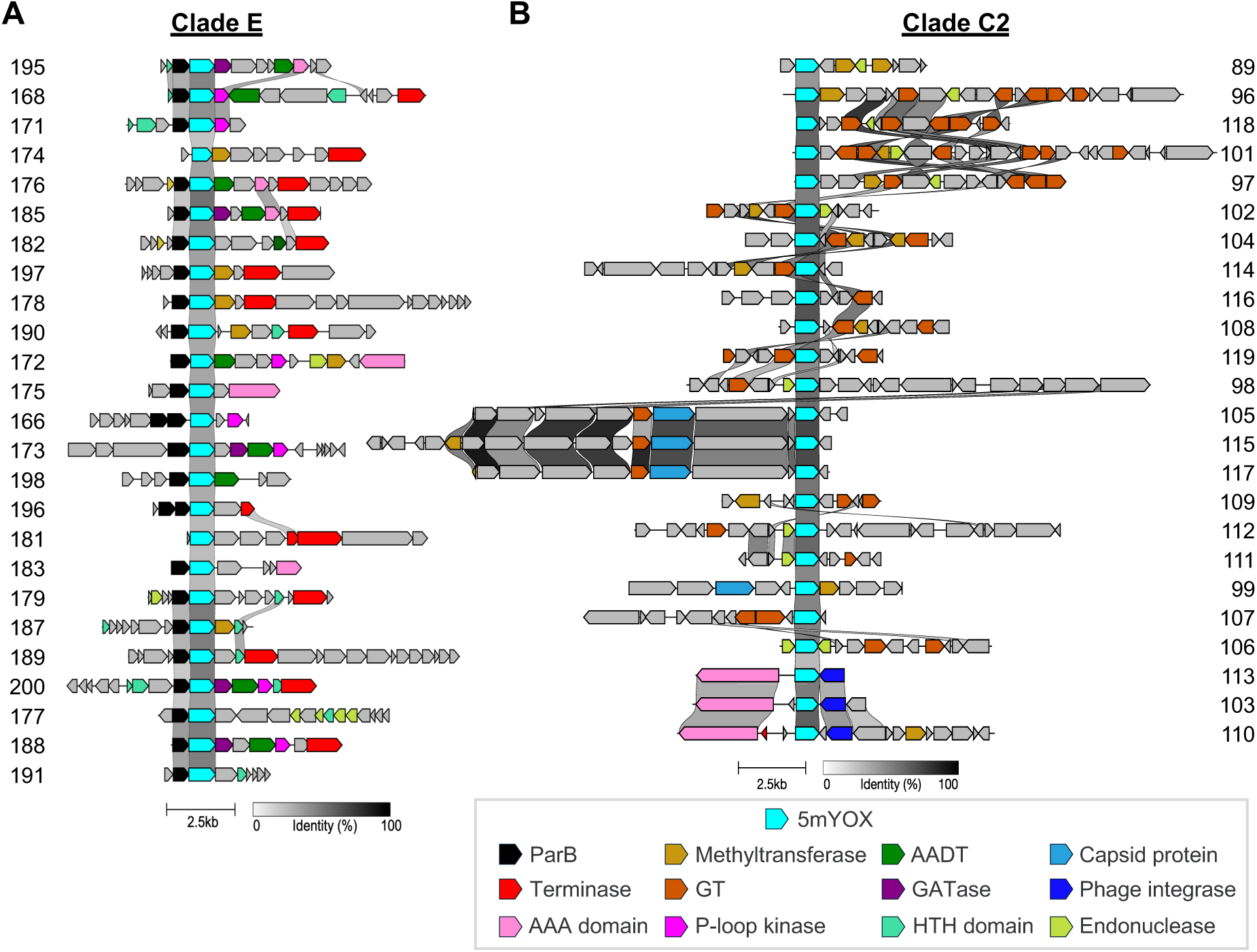
Genomic neighborhoods of Clade C2 and E 5mYOXs. *(A)* Clade E and *(B)* Clade C2 contigs were annotated using MetaGeneMark for gene identification and HMMER3 (50) for domain annotation and/or structure homolog searching via AF2 and Foldseek (Materials and Methods). AAA = triple-A ATPase; HTH = Helix-Turn-Helix.

For experimental characterization, we curated sequences from the SSN clusters that diverged away from trypanosome JBPs (Fig. 2B; *E*-value: 1 × 10^-50^, purple dotted circle; and *E*-value: 1 × 10^-70^, magenta dotted circle) using a multi-step process involving sequence clustering, length filtering, and identity-based reduction (Figs. S1 and S2; Materials and Methods). MSA and phylogenetic analyses show that sequences from the 5mYOX89 cluster (Fig. 2B; purple dotted circle) consistently grouped within the previously classified Clade C2 (7), while Clade C1 members diverged into distinct subclades, including a newly defined Clade E (Fig. S1A).

Analysis of gene architectures of the coding DNA sequences (CDSs) within these subclades revealed that Clade E 5mYOXs co-localize with a homolog of the bacterial chromosome partitioning protein ParB (32, 33) and are consistently linked with known T-hypermodification machinery, such as glutamine amidotransferases (GATases), P-loop kinases, and amino acid:DNA transferases (AADTs, formerly referred to as aGPT-Pplases) (37, 38) (Fig. 3A). Clade C2 enzymes lack ParB and are frequently associated with genes encoding glycosyltransferases (GTs), cytosine or adenosine methyltransferases, and endonucleases (Fig. 3B). Together, these distinctions in SSN, phylogeny, and gene architecture suggest divergent biological roles for Clade C2 and E enzymes.

### High-throughput (HT) functional screening of active T-dioxygenases and their cognate ParBs in *Escherichia coli (E. coli)*

To experimentally assign function to representative 5mYOXs from Clades C2 and E (Figs. S1B and C) and investigate any potential role for a ParB-like partner protein in Clade E activity, we conducted HT *E. coli*-based functional screening using a previously established protocol (23). In brief, *E. coli* was transformed with plasmids encoding either 5mYOX alone or, where appropriate, in combination with ParB. Proteins were co-expressed in a plate format, followed by cell pellet harvesting, genomic DNA extraction, and nucleoside digestion. The conversion of T to 5hmU was then analyzed via ultra-high-performance liquid chromatography (UHPLC) and assemblies were validated by whole plasmid sequencing (WPS) (Fig. 4A).

**Fig. 4.**
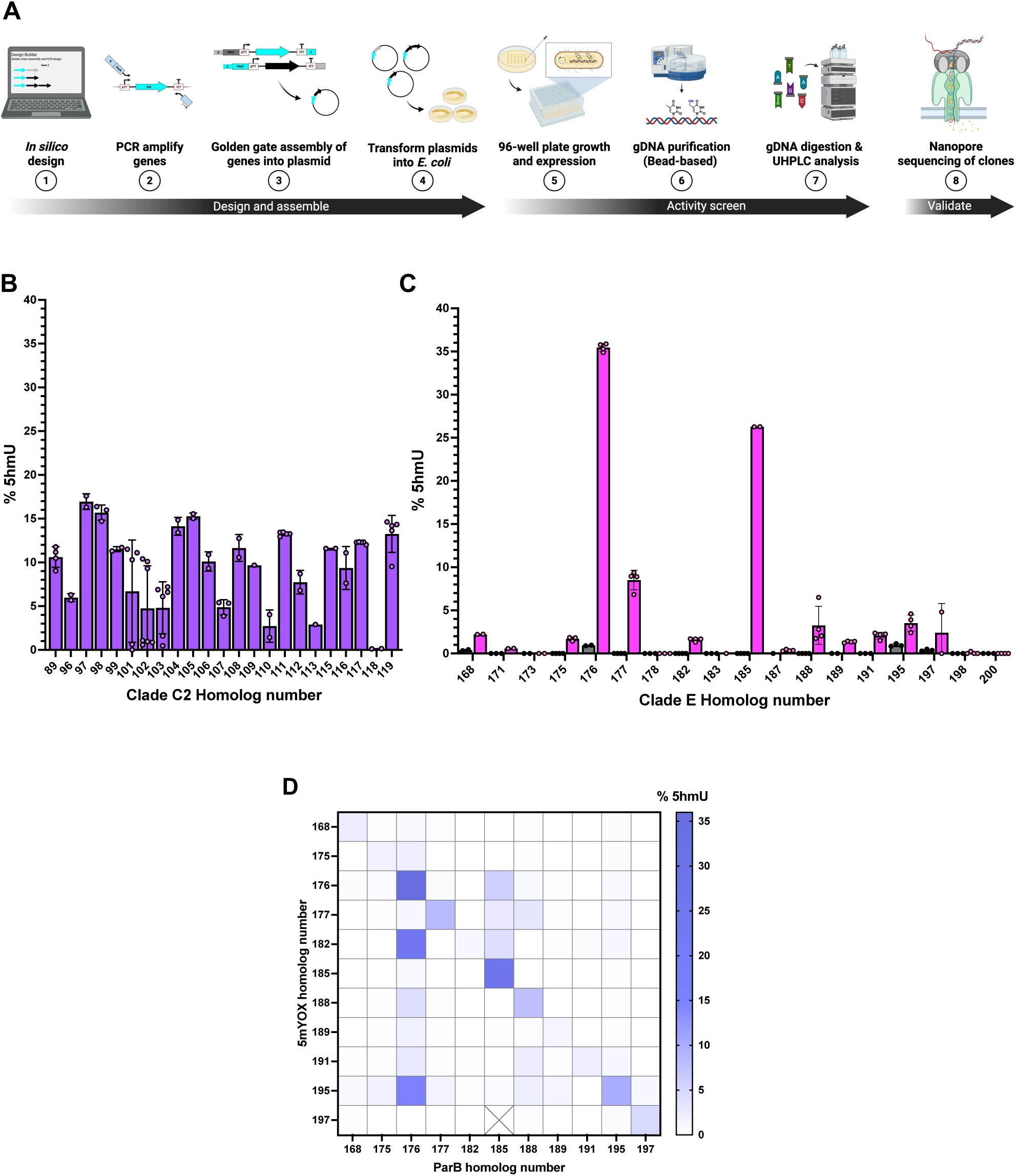
In vivo functional characterization uncovers regulator-dependent and independent T-dioxygenases. *(A)* Schematic of assay platform “Created in BioRender. O’toole, K. (2025) https://BioRender.com/tahrmuf”. UHPLC-based activity of *(B)* Clade C2 and *(C)* Clade E homologs showing the % 5hmU formed on *E. coli* gDNA upon expression of 5mYOX alone or co-expression with ParB. Individual replicates are shown as circles with the error bars representing standard deviation. *(D)* Heatmap showing average % 5hmU on *E. coli* gDNA for non-cognate pairings of Clade E 5mYOX and ParBs. Details are in Tables S1-S3.

Clade C2 homologs were found to hydroxylate T to 5hmU independent of a partner protein, with most members achieving 9-15 % hydroxylation of all T residues in the *E. coli* genome (Fig. 4B and Table S1). Only one member, 5mYOX118, exhibited no activity. In contrast, Clade E 5mYOXs displayed minimal T to 5hmU conversion when expressed alone (0.5-1 %; Fig. 4C and Table S2). However, co-expression of 5mYOX176 and 185 with cognate ParBs significantly enhanced their hydroxylation activity resulting in ∼ 25-35 % 5hmU. Similarly, 5mYOX177 achieved ∼ 8 % hydroxylation in the presence of ParB, highlighting the essential role for ParB co-expression to achieve efficient T hydroxylation by these Clade E 5mYOXs. The remaining Clade E enzymes showed only 0-2 % hydroxylation even in the presence of ParB. ParB alone displayed no activity (Table S2).

To determine whether elevated protein expression contributed to hydroxylation efficiency in these enzymes, we examined 5mYOX/ParB pairs with differing activities. Despite comparable protein levels, 5mYOX/ParB 182 and 198 exhibited substantially lower activity (0.8 and 5 %, respectively) than the ∼ 28 % 5hmU observed for 176. Thus, different expression levels do not account for the observed activity differences (Fig. S3).

Additionally, we assessed the specificity of ParB for its cognate 5mYOX through “mix-and-match” experiments. Different ParBs were paired with 5mYOXs that had previously achieved ∼ 1.5-35 % 5hmU, and T oxidation was monitored (Fig. 4D and Table S3). In some cases, such as with 5mYOX185, enzymatic activity was only observed with its cognate partner. Conversely, 5mYOX182 and 195 demonstrated activity not only with their cognate ParB but also with non-cognate ParBs, showing higher or comparable activity when paired with ParB176. Certain ParBs, including ParB176, 185, and 188 exhibited broad specificity, successfully pairing with 3-9 different non-cognate 5mYOXs. ParB176 also slightly enhanced the oxidation activity of 5mYOX171, 179, and 198 that otherwise showed less than 1 % 5hmU with their native ParBs (Fig. S4A and Table S3).

Although Clade C2 5mYOXs lack a neighboring ParB gene, we tested whether ParBs could enhance their activity by co-expressing ParB176. In most cases, this did not result in a substantial change, except for 5mYOX98 where noticeable decrease in 5hmU levels was observed, while co-expression with 5mYOX107 led to a modest increase (Fig. S4B and Table S3). In vitro examination is required to understand the variation in these two specific cases.

Collectively, these findings indicate two functional subclasses of T-dioxygenases: Clade C2 members that oxidize T independently, and Clade E 5mYOXs that require the co-expression of a ParB-like protein for activity, an observation not previously noted for 5mYOXs.

### MSA and AlphaFold2 (AF2)-guided identification of domains essential for T oxidation

AF2 predictions and Foldseek structural homolog searches were conducted with all Clade C2 and E 5mYOXs to assess structural similarities (39, 40) (Table S4). The top hits for both clades were X-ray crystal structures of human TET2 (hTET2), which within each other exhibit minimal conformational differences (RMSD ≤ 1.5 Å) (11, 41, 42). Accordingly, we aligned AF2-predicted structures to hTET2 PDB: 4NM6 structure bound to 5mC-DNA, NOG, and Fe(II) (11, 39). Using this alignment and MSA, we confirmed conservation of key catalytic residues previously identified in eukaryotic 5mYOXs, and observed two distinguishing features: a “variable insertion” domain positioned between residues 73-118 (consensus sequence) present in both clades, and an extended C-terminus only found in Clade E, which aligns with the C-terminus of hTET2 in the AF2 model (Figs. 5A, B, and S5).

**Fig. 5.**
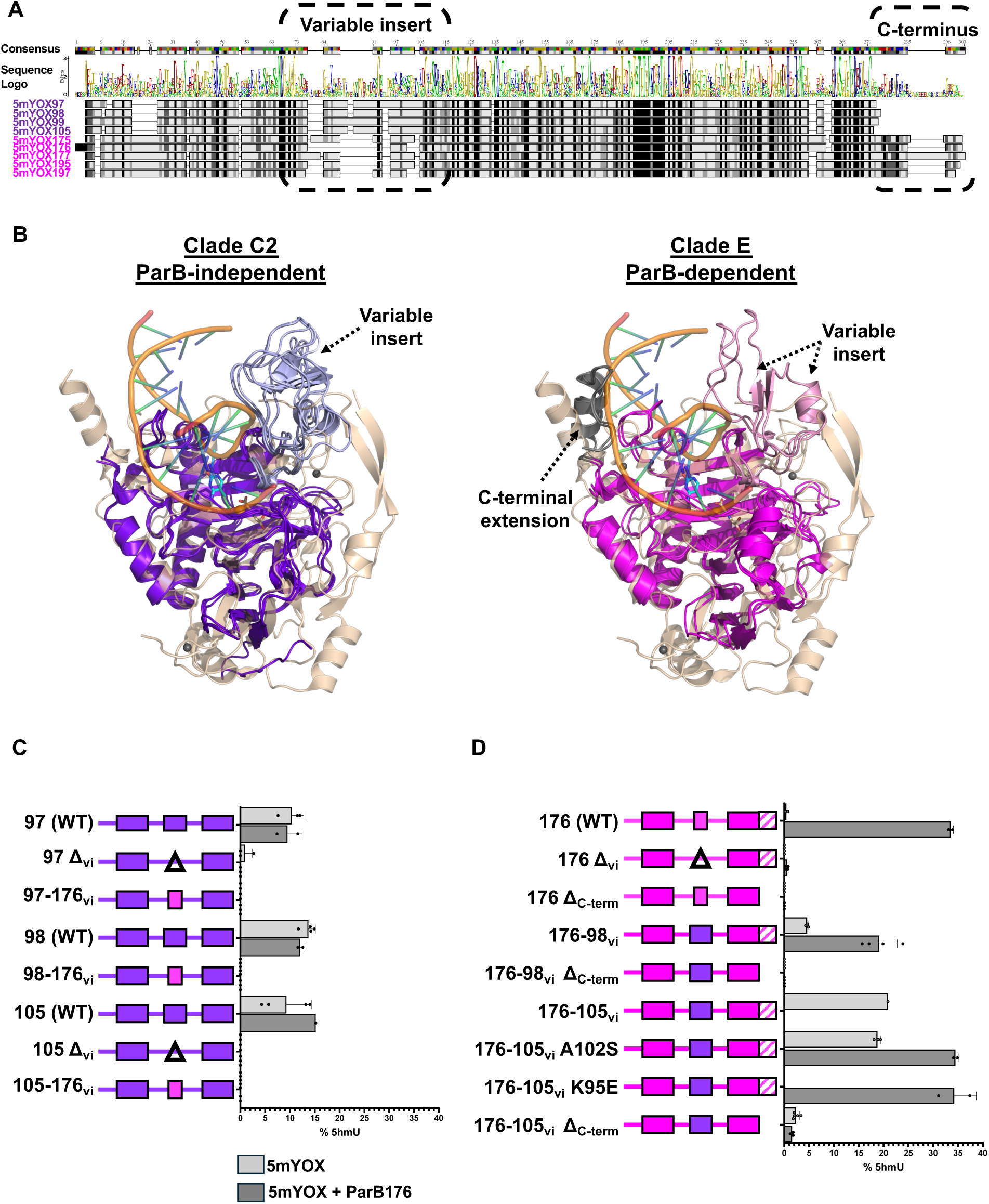
Domains essential for T oxidation. *(A)* MSA of Clade C2 5mYOX 97, 98, 99 105 and Clade E 5mYOX175, 176, 177, 195, 197 generated using the MAFFT algorithm on Geneious Prime. (*B*) Structural overlays of hTET2 (PDB: 4NM6, wheat) and AF2 structural predictions of 5mYOX97, 98, and 105 (Clade C2, purple) or 176 and 195 (Clade E, magenta). 5mC (cyan) and 2OG (wheat) co-substrates are shown as sticks while metals iron (orange) and zinc (gray) are shown as spheres. Variable insertion domain is shown in light purple and light pink for Clade C2 and E, respectively. In Clade C2, this domain is 45-62 residues in length (average 59 residues), compared to 26-52 residues in Clade E (average 43 residues). The C-terminal extension of Clade E is colored gray. AF2 prediction pLDDT and pTM scores are as follows: 5mYOX97 (97, 0.926), 5mYOX98 (93.7, 0.912), 5mYOX105 (91.9, 0.813), 5mYOX176 (94.4, 0.911), and 5mYOX195 (96.4, 0.92). (*C*) UHPLC-based activity data of Clade C2 and (*D*) Clade E variants. Individual replicates are shown as circles with the error bars representing standard deviation. Detailed data in Table S5.

To test whether the variable insertion or C-terminal domains contribute to regulator dependence, we generated domain-deletion and domain-swap variants of select Clades C2 (97, 98, and 105) and E (176) homologs (Figs. 5C, D, and Table S5). The selection was based on the enzymes’ high activity and reproducibility across biological replicates. Interestingly, replacement of the variable insertion domain with a short SG linker in these homologs abolished activity regardless of ParB, demonstrating that this domain is essential for oxidation. When replacing the variable domain in Clade C2 homologs with that of Clade E 5mYOX176, generating a Clade C2– E chimeric protein, no activity was observed (Fig. 5C and Table S5). However, when swapping the variable insertion domain of Clade E 5mYOX176, with that of Clade C2 105 or 98, creating a Clade E–C2 chimeric protein, considerable activity was detected both in the absence and presence of ParB (Fig. 5D and Table S5). Specifically, 5mYOX176-105_vi_ generated 20 % 5hmU without ParB and 30 % 5hmU when ParB was co-expressed. Notably, a mutation in the variable insertion domain, either A102S or K95E, was observed in different biological replicates of 5mYOX176-105_vi_ though the functional relevance of these mutations is unclear. The effect of Clade C2 5mYOX98_vi_ on the activity of 5mYOX176-98_vi_ chimera was not as significant as that of 105_vi_ (4 % 5hmU without ParB and 19 % 5hmU with ParB), but T oxidation was reproducibly detected, regardless of ParB presence. These findings demonstrate that specific Clade C2 variable insertion domains can substitute for the regulatory function of ParB in Clade E 5mYOX176, suggesting that ParB promotes a structural configuration for activity that these domains intrinsically provide.

The C-terminal extension unique to Clade E is also essential for T oxidation. Truncating 5mYOX176 or 5mYOX176-105_vi_ to the last residue conserved across Clades C2 and E (C280) reduced activity to < 1% and 2 %, respectively, regardless of ParB (Fig. 5D and Table S5). This reduction mirrors the inactivity of wild-type 5mYOX176 in the absence of ParB, suggesting that the C-terminal extension contributes to catalytic activity through a ParB-independent mechanism.

### Comparative analysis of bacteriophage 5mYOX-associated ParBs to bacterial ParBs

Through HMM profile-based searching and previous literature reports, we identified and annotated 5mYOX-associated ParBs that match the ParB/sulfiredoxin (Srx) Pfam profile (PF02195) (24, 25). In some contigs, initial annotation was not sufficient to identify ParB-encoding genes, requiring further analysis via AF2/Foldseek and/or structure-based alignments. These analyses consistently reveal *Myxococcus xanthus* ParB (MxParB) (32) and *Caulobacter crescentus* ParB (CcParB) (33) as the closest structural homologs of 5mYOX-associated ParBs (Table S6) and predict that the 5mYOX-associated ParBs contain the N-terminal nucleotide-binding domain (NBD), involved in CTP binding and hydrolysis, and the C-terminal dimerization domain (CTD) (Fig. 6A). However, 5mYOX-associated ParBs appear to lack the DNA binding domain (DBD) present in classical DNA partitioning ParBs. Instead, they contain a ∼ 30-50 amino acid segment that is non-conserved and unstructured according to the AF2 model (Fig. 6A).

**Fig. 6.**
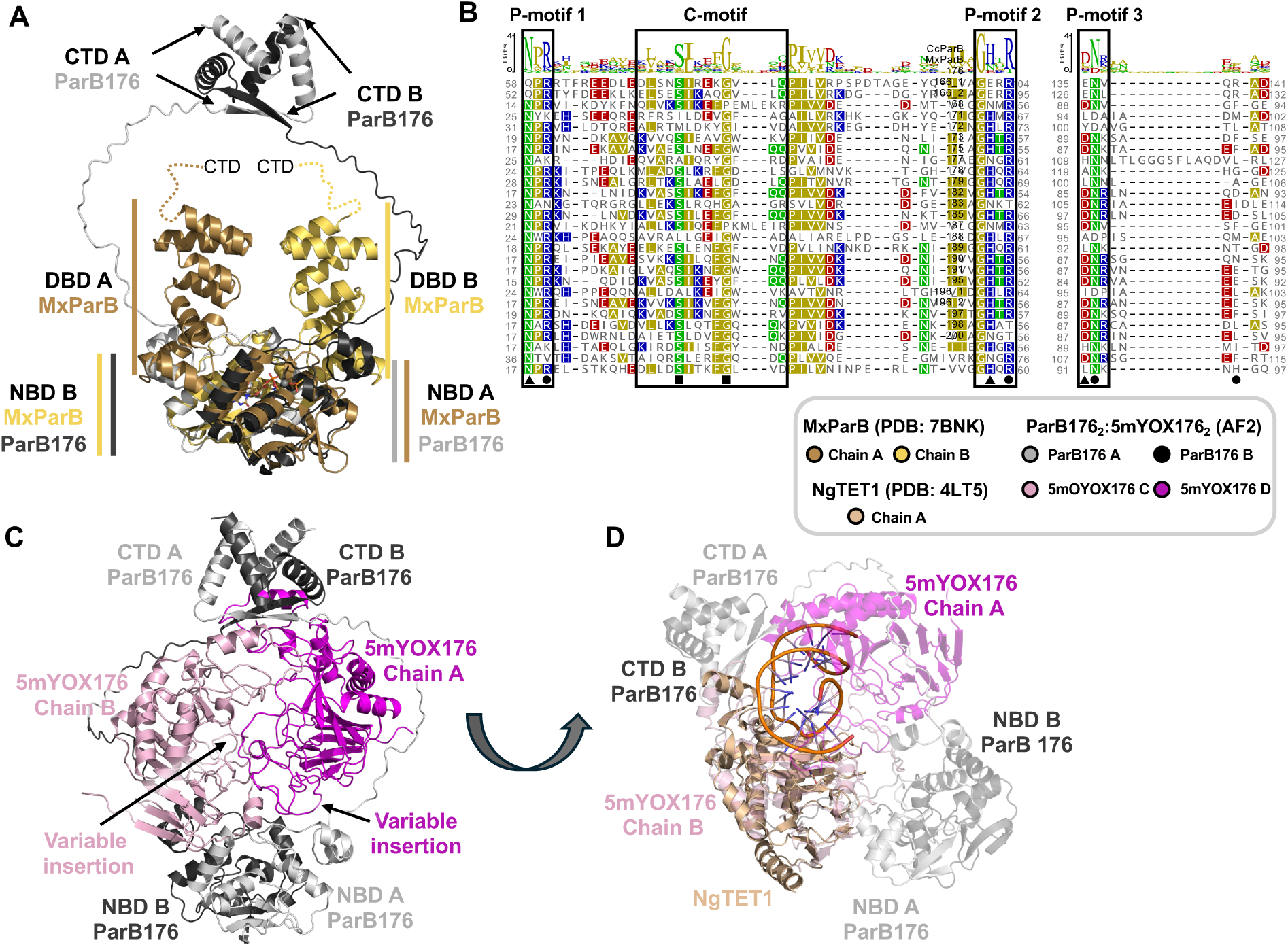
Comparative analysis of 5mYOX-associated ParBs and bacterial DNA partitioning ParBs. *(A)* Structural alignment of ParB176_2_:5mYOX176_2_ AF2 multimer prediction (dark/light gray, pLDDT score 74.7, pTM score 0.467) with MxParB (PDB: 7BNK, gold/yellow) bound to Mg^2+^ (gray sphere), CDP and monothiophosphate (PO_3_S) (sticks). MxParB is a CTD-deletion variant (32). 5mYOX domains in the multimer prediction are hidden to highlight ParB domains. *(B)* MSA of Clade E ParBs, CcParB, and MxParB. Active site residues: triangle-CTP hydrolysis; circle-interactions with CTP phosphoryl groups; square-interactions with cytosine base. Motif nomenclature according to Jalal *et al*. (33). *(C)* AF2 ParB176_2_:5mYOX176_2_ multimer prediction. *(D)* 5mYOX176 (molecule B, light pink) from ParB176_2_:5mYOX176_2_ AF2 multimer aligned with NgTET1 (PDB: 4LT5, wheat) bound to Mn^2+^ (gray sphere), N-oxalylglycine (wheat sticks), and DNA (5mC base flipped into active site).

Due to overall low sequence identity between MxParB, CcParB, and 5mYOX-associated ParBs (∼ 5-14 % ID), we were unable to generate high-confidence MSAs of the CTD. However, alignment of the NBDs was successful and allowed us to compare the conservation of previously identified bacterial key residues (32, 33) in phage ParBs (Fig. 6B and S6). Residues responsible for CTP binding, either to the cytosine base or the phosphate groups, appear to be moderately, though not strictly, conserved between phage and bacterial ParBs. One exception is MxParB arginine (R) 130, which is not conserved in phage ParBs. Residues involved in CTP hydrolysis, including MxParB glutamine (Q52) and glutamic acid residues (E93 and E126), are variably conserved in phage ParBs. Q52 is strictly conserved as asparagine (N) across 5mYOX-associated ParBs. E93 is replaced by either aspartic acid (D) or histidine (H), while E126 aligns with D88 in ParB176; however, this latter position is not conserved across all phage ParBs. Together, these observations suggest that 5mYOX-associated ParBs may retain the structural fold of DNA partitioning ParBs but represent a highly divergent and previously uncharacterized branch of the ParB/Srx structural family. Furthermore, it suggests that 5mYOX-associated ParBs may have evolved distinct functional roles tailored to the regulation of T-dioxygenases, opening new avenues for understanding how structural scaffolds can be repurposed for specialized enzymatic control.

Guided by biochemical and crystallographic evidence of bacterial ParB dimerization (32, 33), we generated an AF2 multimer structure prediction of ParB176_2_:5mYOX176_2_ (Fig. 6C), which predicts the two NBD and CTDs of ParB dimerizing, encapsulating the two 5mYOX domains in the middle (pLDDT score: 74.7). When aligning MxParB (PDB: 7BNK) to the ParB domains of the AF2-predicted complex, the NBD aligns well to the predicted dimer (Fig. 6A) (32). The Predicted Aligned Error (PAE) plots, which reflect the confidence in the predicted distance between two residues, indicate low confidence in the relative positioning of the NBD domains of ParB176 with 5mYOX176 and in the positioning of the two 5mYOX176 subunits with respect to each other (Fig. S7). However, there is reasonable confidence in the proximity of the C-termini of ParB and 5mYOX176 (Figs. 6C and S7). When aligning the X-ray crystal structure of *Naegleria gruberi* TET1 (NgTET1) (PDB: 4LT5, top Foldseek hit among Clade E 5mYOXs, Table S4) to one molecule of 5mYOX176, the DNA is positioned right at the center of the complex, between the two 5mYOX molecules (Fig. 6D) (12). This model aligned to the X-ray crystal structure of MxParB shows the 5mYOX molecule in the location that the DNA binding domain of bacterial ParBs is positioned (Fig. 6A and 6C). Although a model, this illustrates a hypothesis that ParB may serve to promote conformational selection of 5mYOX:DNA interactions to allow for T oxidation.

Within the 5mYOX-associated ParBs, phylogenetic relatedness appears to be a determinant of their ability to activate T oxidation (Fig. S8A). All ParBs that show successful activation of 5mYOX (> 1.5 % 5hmU produced) fall within “Clade 1” of the phylogenetic tree, except for ParB175, which belongs to “Clade 3”. In contrast, ParBs from “Clade 2” fail to support this activity. For example, only ParB198 shows minimal activation of its cognate 5mYOX (< 0.1 % 5hmU). Remarkably, all “Clade 2” ParBs lack conservation of at least one major CTP-binding residue (Fig. S8A, circles), which might suggest a mechanistic divergence in activation. These ParBs may require a non-canonical NTP for activation and given that our in vivo assay is constrained by the metabolite composition of *E. coli*, such activity may not be detectable under these conditions.

ParB176, which exhibits promiscuous activity with the ability to activate several non-cognate 5mYOXs (Fig. 4D and Table S3), does so with 5mYOXs that have cognate ParBs within “Clade 1”, except for ParB175 (Clade 3) and ParB198 (Clade 2). Among these, 5mYOX182 exhibits significantly higher 5hmU levels when co-expressed with ParB176 compared to its cognate partner. Of note, ParB182 shares the highest sequence identity with ParB176 (46 % ID), and 5mYOX182 also shares notable identity with 5mYOX176 (50 % ID) (Fig. S8B). Protein expression levels are not a factor in this substantial increase (Fig. S3B). ParB188 is another example that supports this activation trend with successful pairing with 5mYOX177, 5mYOX191, and 5mYOX195 (Fig. S8). In general, if ParB-5mYOX non-cognate pairings are successful to activate T oxidation in vivo, it is most likely that the homologs fall within the same sub-clade in the respective phylogenetic trees illustrating the possible evolutionary basis for interaction specificity between these two classes of proteins.

### Surveying for unexplored 5mYOX pathways

With two newly defined subclasses of phage-encoded 5mYOXs in hand, we surveyed publicly accessible databases to assess the prevalence and diversity of these pathways in nature. Using HMM profiles generated for Clade C2 (ParB-independent T-dioxyggenases), Clade E (ParB-dependent T-dioxygenases), and 5mC-dioxygenases (Clades A/B) (7), hereto referred to as the phage-5mYOX HMM profile, we searched a broad range of archaeal, viral, and bacterial genome databases (Materials and Methods), focusing on organisms known to assemble genes in operons/biosynthetic gene clusters (Fig. 7 and Table S7).

**Fig. 7.**
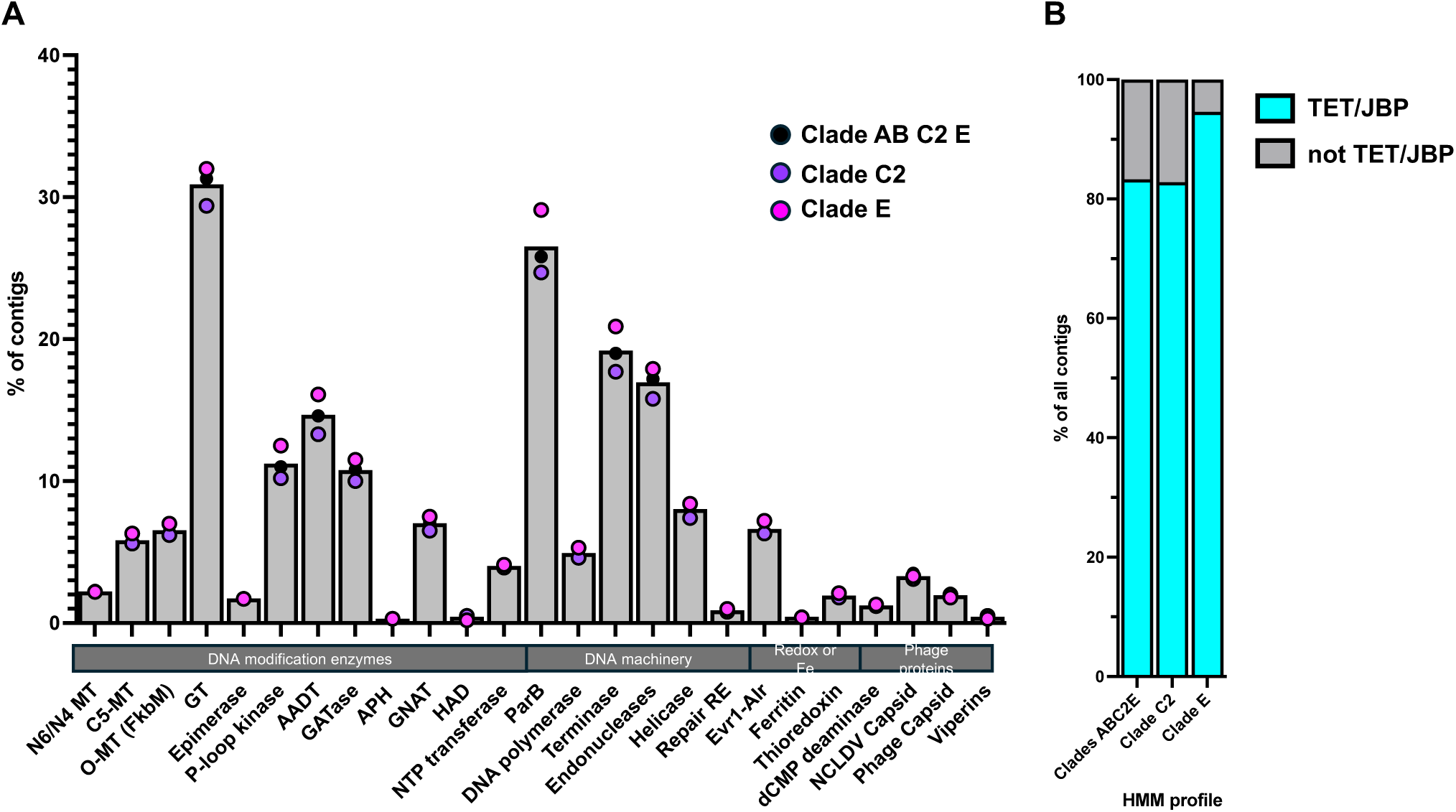
Genome mining of metagenomic databases using custom HMM profiles reveals DNA hypermodification machinery. *(A)* Summary of domain annotations in contigs mined using Clades ABC2E (black), Clade C2 (purple), or Clade E (magenta) HMM profiles. Gray bar represents the average % of contigs from all three searches containing each domain annotation. N6/N4 MT = N6-adenosine/N4-cytosine methyl transferase; O-MT = O-methyltransferase; APH = aminoglycoside 3’-phosphotransferase; GNAT = Gcn5-related N-Acetyltransferase; HAD = haloalkanoate dehalogenase; RE = restriction enzyme; NCLVD = nucleocytoplasmic large DNA viruses. See Table S7 for full genome mining and Pfam data. *(B)* Summary of contigs from each HMM profile: Clades ABC2E (phage-encoded 5mC- and T-dioxygenases), Clades C2 (T-dioxygenase, ParB-independent), and Clade E (T-dioxygenase, ParB-dependent) that co-occur with TET/JBP Pfam annotations.

Across all searches, the most frequently co-occuring domains were GTs (31 %), ParBs (27 %), terminases (19.2 %), and endonucleases (17 %). In addition, we identified several different classes of DNA hypermodification genes in contigs including AADTs, P-loop kinases, NTP-transferases, sugar epimerases, O-methyltransferases, and GATases (Fig. 7A and Table S7). Strikingly, only 6 % of all pathways co-occur with a C5-MT, suggesting that the majority of all 5mYOXs mined (94 %) produce 5hmU rather than 5hmC. Additional genes such as helicases (8 %), DNA polymerases (5 %), and iron-sulfur maturation protein Evr1-Alr (7 %) were also found in a subset of contigs.

Notably, while most enzyme families were similarly distributed across hits from all HMM profiles, the Clade E HMM profile retrieved 4-5 % more ParB-containing contigs compared to the Clade C2 or phage-5mYOX HMM profiles (Fig. 7B and Table S7), consistent with its functional dependence on this partner protein. Moreover, the Clade E 5mYOX HMM profile showed the strongest overlap with the TET/JBP Pfam HMM profile (PF12851), with 94.6 % of all retrieved contigs containing this annotation, compared to 83 % of contigs retrieved using either the Clade C2 or phage-5mYOX profiles. Remarkably, thousands of contigs lacked a TET/JBP annotation altogether, indicating either that they may represent highly divergent members of this enzyme family and/or that the phage-specific 5mYOX HMM profiles generated in this study may better represent 5mYOXs present in these databases.

Together, these findings reveal that 5mYOX pathways are not only widespread but also remarkably diverse in their genomic contexts. The discovery of thousands of unannotated, potentially novel 5mYOX-like sequences hints at a vast, untapped reservoir of DNA-modifying enzymes in nature.

## Discussion

The 5mYOX superfamily, typified by its Fe(II)/2OG-dependent dioxygenase activity, is prevalent across all domains of life. Until recently, nearly all functional insight has come from eukaryotic representatives, particularly those involved in 5mC oxidation. Our previous work was the first to demonstrate that bacteriophages encode functional 5mC-dioxygenases (7), expanding the enzymatic landscape of this family. In this study, we present evidence for T-dioxygenases of phage-origin, representing the first comprehensive study of T-dioxygenases outside of eukaryotes (Figs 2-4). These phage-derived enzymes, while sharing the core structural fold and catalytic residues of the broader 5mYOX family (Figs. 5 and S5), are distant in sequence and exhibit remarkable diversity.

Through HT functional screening of ∼ 50 phage-derived 5mYOX homologs, we identified two functionally distinct subclasses: Clade C2 enzymes, which catalyze T oxidation independently, and Clade E enzymes, which require co-expression of a ParB-like regulator for T oxidation (Figs 2, 3, and S1). This discovery represents a previously unrecognized regulatory split within the 5mYOX family and introduces a novel paradigm for dioxygenase activation. To probe the molecular basis of this functional divergence, we performed domain-swapping experiments that revealed the variable insertion domain of Clade C2 is sufficient to confer partial or full ParB independence when introduced into a Clade E backbone. Strikingly, the resulting chimera, 5mYOX176-105_vi_ (Clade E–C2 chimera) retained robust activity in the absence of ParB and was further stimulated by ParB co-expression, underscoring the modular and tunable nature of these domains. In contrast, the reverse chimera (Clade C2–E chimera) failed to restore activity even when co-expressed with the corresponding ParB, suggesting that additional structural features or context-specific interactions are required to support catalysis.

Additionally, deletion of a C-terminal extension domain specific to Clade E impaired the activity of both 5mYOX176:ParB176 and the ParB-independent chimera 5mYOX176–105_vi_, strongly implicating it in the oxidation function (Figs. 5C and D). Structural overlays show that this region aligns closely with residues 1909–1924 of hTET2 (Figs. 5B and S5), and prior studies have demonstrated that deletions beyond residue 1913 significantly compromise hTET2 catalytic activity (11). These findings suggest that this motif may represent a structural feature essential for maintaining dioxygenase function in some dioxygenase classes. Intriguingly, AF2 multimer modeling of the 5mYOX176₂:ParB176₂ complex predicts this same C-terminal domain to be positioned at the dimer interface of ParB and 5mYOX (Fig. 6C), suggesting that this domain acts both as a catalytic determinant and a regulatory interface that enables dynamic tuning of enzyme activity.

5mYOX-associated ParBs retain the characteristic NBD and CTD, but lack the DBD, of their distant homologs in bacteria (Fig. 6A). In the bacterial ParBs, the NBD binds to CTP triggering dimerization of this protein and the formation of a clamp-like structure, which allows its sliding along the DNA, dissociating upon CTP hydrolysis (32, 33). In our system, 5mYOX-associated ParBs can stimulate T oxidation in *E. coli* (Figs. 4C-D and S8), and structural predictions position the 5mYOX as the DNA binding domain (Fig. 6D) with two phage 5mYOX monomers brought into proximity upon dimerization of the bound ParB molecules to function on DNA (Figs. 6C-D, and S7). This raises the possibility of a higher-order 5mYOX-ParB complex that coordinates oxidation events, potentially mirroring the assembly dynamics of the ParABS system.

The physiological role of the 5hmU modifications resulting from 5mYOX oxidation of T in bacteriophage remains an open question. Expression of phage-encoded T-dioxygenases in *E. coli* only results in partial oxidation of genomic T sites (∼ 10-20 %) and the variability in 5hmU levels across homologs suggests potential sequence-specificity patterns. Additionally, our comprehensive genome mining reveals hypermodification machinery associated with T-dioxygenases that could be involved in glycosylation, O-methylation, and amino acid functionalization (Figs 3, 8, and Table S7). Several phage 5mYOX pathways mined co-occur with endonucleases which could suggest methods in which the virus targets its bacterial host while protecting its own DNA via base modification. Furthermore, the vast majority of contigs mined are predicted to be T-hypermodification pathways, based on the absence of an annotated C5-MT (Pfam PF00145) in the nearby genome. The enrichment of putative T-dioxygenase pathways over 5mC-dioxygenases in our mining results may reflect a broader biological trend in which thymine serves as a more versatile substrate for viral genome adaptation, as 5mC modification is more linked to host regulatory pathways. In fact, the co-occurrence of ParB with Clade E 5mYOXs could serve to regulate timing of 5mYOX activation in the life cycle of the virus, and/or conformational control of the cognate 5mYOX to selectively bind certain T sites on DNA and enabling oxidation.

All in all, this study uncovers a previously uncharacterized class of the 5mYOX superfamily, reveals a novel mode of enzymatic regulation via ParB-like proteins, and opens the door to a vast and largely unexplored landscape of T hypermodification pathways in phage biology. These findings also lay the groundwork for future exploration into the biochemical and evolutionary significance of DNA modification in viral systems.

## Supporting information

Table S1. Clade C2 in vivo activity data.

Table S2. Clade E in vivo activity data.

Table S3. Non-cognate 5mYOX-ParB in vivo activity data.

Table S4. Summary of Foldseek results for phage-encoded 5mYOX enzymes.

Table S5. Chimeric 5mYOX in vivo activity data.

Table S6. Summary of Foldseek results for phage-encoded ParB proteins.

Table S7. Genome mining statistics.

Table S8. Scalar normalization values for relative quantification of 5hmU by UHPLC peak areas.

## Author contributions

K.H.O. and L.S. designed research; K.H.O., A.B., M.L.D. and S.R.L. performed research; K.H.O., A.B., M.L.D., S. R. L., S.B., H.B., and L.S. contributed new reagents/analytic tools; K.H.O., A.B., M.L.D., S.R.L., and L.S. analyzed data; K.H.O and L.S. wrote the paper.

## Acknowledgements

We appreciate the support of Drs. Zhiyi Sun, Sean Johnson, Peter Weigele, and Vladimir Potapov on bioinformatic pipeline development and infrastructure, and Dr. Nan Dai with analytical methodology. We also thank Dr. Jackson Buss and Mr. Ray Moncion for optimized PCR and plasmid DNA purification protocols and the NEB sequencing core for both Sanger and WPS of constructs and plasmid assembly verification. We are especially grateful for continued discussions and insight from Drs. Sean Johnson and Peter Weigele and careful review of the manuscript from Drs. Tim Blower and Rebekah Silva. This work was supported by funding from New England Biolabs.

## Materials and Methods

### Chemicals, media, and cell strains

Unless otherwise noted, all chemicals are obtained from Millipore Sigma and used without additional purification. Media and media components were sourced from Beckton Dickinson-Difco. All competent cells and cloning reagents or kits were from NEB. Plasmids and genes were sourced and synthesized via Twist and Genscript as described in a following section. DNA purification kits were obtained from NEB unless otherwise specified.

### Sequence dataset curation and SSN generation

Sequence dataset curation for SSN generation (Fig. 2) included sequences from UniProt and from JGI IMG/VR v3 (36). All sequences in the following InterPro/Pfam families were included: PF12851 (TET/JBP), IPR024779 (2OGFeDO, oxygenase domain), and IPR040175 (Methylcytosine dioxygenase TET1/2/3) (34, 35). Sequences from JGI IMG/VR 3 database were mined using HMM profile for TET/JBP superfamily (PF12851) (35, 36). Additional sequences were manually curated based on literature predictions (28) and further curated using literature sequences as seed sequences to search against NCBI. All-by-all Basic Local Alignment Search Tool (BLAST) calculations were run using the EFI-EST (https://efi.igb.illinois.edu/efi-est/) (30, 31) with alignment score thresholds of 35, 50, and 70 using network connectivity scoring. The alignment threshold and representative node percentage were selected based on empirical analysis. Of the 10,897 sequences, 6,381 are nonredundant sequences. Networks are visualized using Cytoscape (43). To extract homologs from seed sequences, we employed networks at two different alignment score thresholds (50 and 70), based on the enzyme characteristic we were probing.

For curating homologs of seed sequence “5mYOX89”, a 40 % representative node network with an alignment score of 50 from sequences represented in Fig. 2B, purple-dotted circle was used. The cluster containing 5mYOX89 was extracted (86 sequences), and then further filtered using CD-HIT filtering by 75 % identity resulted in 20 sequences (44, 45). Four additional sequences were manually selected, resulting in a sequence dataset of 24 homologs between 24 and 77 % identical (excluding 5mYOX89) (Fig. S1A (Clade C2) and S1B).

The cluster containing Clade C1 homologs in the 1 × 10^-70^ network was selected for homolog prioritization (Fig. 2B). The cluster was extracted (278 representative sequences), and then further filtered by sequence length, removing all sequences under 160 amino acids (260 sequences), followed by another round of filtering by 70 % identity using CD-HIT (121 sequences) (44, 45). Representatives of this sequence set were manually curated based on phylogenetic tree distribution, prioritizing a few homologs from each colored clade as shown in Figs. S1A (Clade C1) and S2. This resulted in a diverse and representative sequence set of 34 sequences that are between 26 and 62 % identical (Fig. S1A (Clade E) and S1C).

### Cloning of individual genes into destination plasmid

Gene fragments were ordered from Twist Bioscience (South San Francisco, CA) for Clade C2 homologs (5mYOX96-119) containing the appropriate overhangs for GGA (NEB BsaI HFv2 (R3733S, NEB, Ipswich, MA) and NEB T4 DNA ligase (M0202T, NEB, Ipswich, MA)), codon optimized for *E. coli* expression and domesticated to remove any BsaI restriction enzyme cut sites. GGA was conducted into destination plasmid pSL003 as described previously (23). The 3′-end of the open reading frame defined for 5mYOX108 was not successfully sequenced and therefore resulted in 9 undefined amino acids at C-terminus. We decided to order the gene truncated back to the last conserved residue across Clade C2 5mYOXs (5mYOX108 Cys290).

Clade E homolog genes (5mYOX and ParB) were codon optimized for *E. coli* expression, domesticated to remove any BsaI restriction enzyme cut sites, and synthesized from oligo pools ordered from Twist Bioscience (South San Francisco, CA) as described by Lund *et al.* (46). Synthesized genes, also containing appropriate overhangs for GGA, were assembled into pSL003 destination plasmid as described previously (23). The following gene fragments were synthesized by GenScript (Piscataway, NJ) and assembled following the same parameters described above: ParB166_1, ParB172, ParB173, ParB178, ParB185, 5mYOX181, and 5mYOX187.

Chimeric enzyme genes were ordered as gene fragments from GenScript (Piscataway, NJ) with the appropriate overhangs for GGA, codon optimized for *E. coli* expression and domesticated to remove any BsaI restriction enzyme cut sites. GGA was conducted into destination plasmid pSL003 as described previously (23). Design of gene constructs for variable insert deletion variants was guided via AF2 structure prediction of deletion variants with different length “SG” linkers to predict best possible linker length.

Genes coding for the C-terminal truncation and insertion domain deletion variants of Clade E homologs were cloned from the original or chimeric 5mYOX genes. Primers were designed and PCR conditions were determined following online tools NEBase Changer (https://nebasechanger.neb.com/, NEB, Ipswich, MA) and purchased from IDT (Newark, NJ). PCR reactions were then treated with KLD reaction mixture following the product protocol (NEB, Ipswich, MA) and transformed 5µL KLD reaction into 25 µL NEB 10-beta competent cells (C3019, NEB, Ipswich, MA) in 96-well plate format following product protocol. Individual colonies were selected to inoculate rich media (1.0 % soy peptone, 0.5 % yeast extract, 0.5 % NaCl, 0.1 % MgCl_2_ hexahydrate, 0.1 % dextrose and 50 μg/mL of kanamycin) in a 96-well plate format and grown overnight at 37 °C shaking. Cells were harvested via centrifugation at 2,500 x g for 10 min. The supernatant was decanted, and cell pellets were stored at −20 °C until plasmid purification and sequence verification via WPS described in section below.

Sequence datasets of screened protein sequences for all tested proteins are provided in the supplementary materials.

### BGC design and assembly for pathway expression in *E. coli*

A custom web application “Design Builder” was written for the rapid assembly of many combinations of genes to be expressed in *E. coli*. Briefly, the software acts as an interface for design and selection of assemblies as well as script generator for OT-2 liquid handling robot (Opentrons, Brooklyn, NY).

Users can search for and select individual genes and then generate combinations of these genes in various ways (random, pseudorandom, etc). The application generates two corresponding CSV files, which are subsequently converted to python scripts for use on OT-2 robot (Opentrons, Brooklyn, NY). The first CSV file is used for amplification of individual genes to be used in constructs. Once the PCR has been performed, the resulting concentrations of each gene are uploaded to the software to generate the second CSV file, which instructs the OT-2 in performing the GGA of each construct design using a cherry-picking algorithm. Previously generated design schemes can also be uploaded to the application and modified for additional experimentation.

Assembly of 2 or more genes into a pET28a-based plasmid lacking T7 promotor and terminator (pSL001) using universal primers and GGA was initially described by Pyle *et al*. (23). Briefly, genes were amplified from a pET28a based destination plasmid (pSL003) using primers annealing to T7 promotor and terminator, with a linker and BsaI site. For leave-one-out controls, genes were replaced with pET28a multicloning site inserts. PCR parts were run on a gel to confirm size, and concentration was quantified using Quant-iT Broad-Range dsDNA Assay Kit (ThermoFisher Scientific, Waltham, MA) in a 384-well plate and measured in a SpectraMax Microplate Reader (Molecular Devices, San Jose, CA). PCR part clean-up was performed by adding a 2:1 ratio of sample to plasmid neutralization buffer (T1013 NEB, Ipswich, MA) until homogeneously yellow followed by applying 1.5 x sample volume of SeraSil-Mag^TM^ 400 beads (Cytiva, Marlborough, MA) and 100 % ethanol (1.5:1 Ethanol to beads ratio). PCR parts were then purified using a KingFisher Flex (ThermoFisher Scientific, Waltham, MA) via two 300 µL 80 % ethanol washes, followed by elution in 100 µL Milli-Q water in KingFisher standard plate (ThermoFisher Scientific, Waltham, MA). PCR parts were quantified in 384-well plates using a SpectraMax Microplate Reader (Molecular Devices, San Jose, CA) with a Quant-iT™ Broad-Range dsDNA Assay Kit (ThermoFisher Scientific, Waltham, MA) and normalized to 15 nM using OT-2 liquid handling robot (Opentrons, Brooklyn, NY). GGA reactions were prepared using OT-2, keeping 1:1 molar ratio of all DNA fragments (1.1 nM of each fragment added) and 10 ng destination plasmid (pSL001). GGA reactions were prepared and incubated following the NEB GGA BsaI-HFv2 protocol for multiple gene assembly (NEB, Ipswich, MA) using NEB BsaI HFv2 (R3733S, NEB, Ipswich, MA) and NEB T4 DNA ligase (M0202T, NEB, Ipswich, MA). Assembled plasmids (5 µL) were transformed into T7-express cells (25 µL, C2566, NEB, Ipswich, MA) in deep-well plate format following the standard protocol with a few modifications described here. After heat-shock, recovery incubation occurred in a thermomixer set at 30 °C shaking for 75 min. Cells were pelleted via centrifugation at 2500 x g for 10 min. The supernatant was poured off and pellets were resuspended in remaining volume. Cells were then plated on rich agar (1.0 % soy peptone, 0.5 % yeast extract, 0.5 % NaCl, 0.1 % MgCl_2_ hexahydrate, 0.1 % dextrose, 1.5 % agar) containing 50 µg/mL kanamycin and incubated overnight at 30 °C.

### HT *E. coli in vivo* assay of 5mYOX

Individual colonies were used to inoculate overnight culture in rich media (1.0 % soy peptone, 0.5 % yeast extract, 0.5 % NaCl, 0.1 % MgCl_2_ hexahydrate, 0.1 % dextrose and 50 μg/mL of kanamycin) in 96-well deep well plates (500 µL/well) and grown overnight at 30 °C shaking (250 RPM). Overnight culture was then used to inoculate a 96-well plate (500 μL, 1:100 dilution) containing fresh rich media supplemented with kanamycin (50 µg/mL). Remaining overnight culture was pelleted via centrifugation at 2500 x g for 10 min. Supernatant was decanted, and pellets were either stored at −20 °C or proceeded directly into plasmid purification as described in a subsequent section. Inoculated subcultures were grown at 30 °C for 5 h before inducing with IPTG (400 µM) and grown overnight at 18 °C (16-18 h). Cells were harvested via centrifugation at 2500 x g for 10 min and stored at −20 °C or proceeded directly into gDNA purification. Modified gDNA was purified from *E. coli* as described by Pyle *et al*. (23). Purified gDNA (20 µL) was then digested on bead using nucleoside digestion mix (NEB M0649) overnight at 37 °C as described previously (23).

### UHPLC analysis of nucleosides

Prior to injection on UHPLC, samples were filtered using AcroPrep Advance 0.2 µm plate filters (1200 x g for 10 min) (Product #8019 Cytiva, Marlborough, MA). Nucleosides were subjected to UHPLC on an Agilent 1290 Infinity II UHPLC system (Agilent, Santa Clara, CA) and resolved on a Waters XSelect HSS T3 C18 column (2.1 × 100 mm, 2.5 µm particle size) (Waters, Milford, MA) with a gradient mobile phase consisting of aqueous ammonium acetate (10 mM, pH 4.5) and methanol. Canonical bases were quantified using a scalar approach as described previously (23) and 5hmU was quantified using the same scalar approach, wherein SP8 phage genomic DNA (T completely replaced with 5hmU) was injected onto UHPLC (from 0.1 to 0.4 µg) for scalar calculation (Table S8). The scalar values were then used to normalize UHPLC peak quantification for canonical bases and 5hmU as described previously (23).

### Plasmid purification and Whole Plasmid Sequencing (WPS)

Plasmid purification in plate format was adapted from Monarch Plasmid MiniPrep Kit (T1010, now T1110, NEB, Ipswich, MA) protocol and buffers, with purification using SeraSil-Mag^TM^ 400 beads (Cytiva, Marlborough, MA) in place of columns. Pelleted cells were resuspended in Buffer B1 (150 µL/500 µL culture) and mixed using Thermomixer (1200 RPM for 1 min at 25 °C). Buffer B2 was added (150 µL/500 µL culture) and mixed again using Thermomixer (1200 RPM for 20 min at 25 °C). Buffer B3 (300 µL) was added followed by mixing via Thermomixer until color change observed and homogeneous (1200 RPM for ∼ 30-60 s at 25 °C). Plate was centrifuged for 10 min at 3500 RPM and then lysate (400 µL) was transferred to a KingFisher Deep well plate (ThermoFisher Scientific, Waltham, MA). SeraSil-Mag^TM^ 400 beads (Cytiva Marlborough, MA) were then applied to lysate (425 µL of 2.5 mL bead: 46 mL 100 % Ethanol) and plasmid was purified using KingFisher Flex (ThermoFisher Scientific, Waltham, MA), washing twice with 80 % Ethanol (500 µL) followed by elution in Milli-Q water (100 µL) into KingFisher standard 96-well plates (ThermoFisher Scientific, Waltham, MA).

Concentrations of plasmid were determined using Quant-iT Broad-Range dsDNA Assay Kit (ThermoFisher Scientific, Waltham, MA) as described above. Purified plasmid was then submitted for WPS using Oxford Nanopore sequencing via NEB sequencing core facility. Correct assemblies were verified either using Geneious Prime (Dotmatics, Boston, MA) or using a python script to confirm identity and coverage of genes assembled.

### Protein expression testing

Constructs were assembled and transformed as described above. Overnight cultures were prepared in 96-well deep well plates (500 µL/well) followed by cell pelleting, plasmid purification, and WPS analysis as described above. Prior to cell pelleting, overnight culture was used to inoculate either in a test tube (2 mL, 1:100 dilution, Fig. S3A) or in a 96-well plate (500 µL, 1:100 dilution, Fig. S3B). Growth and induction of protein expression followed by gDNA purification and UHPLC analysis as described above. After protein induction, an aliquot of cells (50 µL) was removed for SDS-PAGE analysis prior to pelleting the remaining culture. Pelleted cells for SDS-PAGE analysis were then resuspended in 1X Blue Protein Loading Dye (B7703S, 30 µL) and run on a Novex^TM^ 10-20 %, Tris-Glycine Plus WedgeWell^TM^ gel (Invitrogen, Waltham, MA). Gels were stained using SimplyBlue^TM^ SafeStain (Invitrogen, Waltham, MA) and destained in deionized water.

### Programs, tools, and bioinformatic analyses

Tools, including ChatGPT (OpenAI) and Microsoft 365 Copilot, were used to assist with language editing and grammar refinement during manuscript preparation. All content were reviewed and verified by the authors for intellectual integrity.

For bacteriophage pathway annotations, contigs were identified using IMG/VR IDs extracted from SSN cluster, and CDS initially defined using Prokka (47) or Geneious Prime software (Dotmatics, Boston, MA). These gene sequences were then extracted and synthesized as described above. The CDS were then annotated for putative function using domainator (48) and Pfam HMM profiles (35). Visualization and analysis of contigs was conducted using Clinker (49) and Geneious Prime (Dotmatics, Boston, MA). Colored domains were annotated by the following Pfam profiles: TET/JBP (PF12851), ParB (PF02195), Terminase (PF20901, PF03237, PF04466, PF17288, PF17289, PF05876, PF05119, PF20441), AAA domain (PF13555, PF00004, PF07728, PF13238), methyltransferase (PF00145, PF01555, PF13651, PF13578, TIGR01177, PF05050), glycosyl transferase (PF00535, PF01755, PF06002, PF05704, PF04488, and CAZY families: GT25, GT45, and GT73), P-loop kinase (PF18751), AADT (PF18724), GATase (PF13522, PF13537), HTH domains (PF01381, PF13730, PF08279, PF20432), capsid protein (PF16903), phage integrase (PF00589), and endonuclease (PF01844, PF01541, PF02945, PF13384, PF16784, PF02796, PF13730, PF08774, PF16784).

Genome mining efforts were further conducted using three unique HMM profiles (1) for bacteriophage 5mYOX broadly (contains Clades A, B, C2, and E 5mYOXs), (2) Clade C2 5mYOXs, and (3) Clade E 5mYOXs. HMM profiles were generated using HMM build (50) from MSAs generated in Geneious Prime (Dotmatics, Boston, MA) using Fast Fourier Transform (MAFFT) with a JTT200 scoring matrix (51). Genomes were mined using the described HMM profiles as query with an *E-*value cutoff of 1 × 10^-5^, Z value of 1000, and deduplicating contigs that are 100 % identical via CD-HIT (44, 45). 10 CDS upstream and downstream of the query hit were extracted. Genome databases searched include: NCBI Representative Genomes (52), IMG/VR 4 (53), wastewater treatment plant metagenome database (54), IMG/M ecosystem metagenomes (permafrost soils, permafrost wetlands, aqua saline, hypersaline, compost, alkaline, hot, near-boiling, root) (55), MetaGPA selected and unselected databases (56), ATCC enterics (ATCC, Manassas, Virginia), Gut phage database (57), Genbank plasmid database (58), IMG/Plasmid resources database (59), RNA Viruses in Metatranscriptomes (60), Thermobase genomes (52), NEB in-house strains (REBASE) (61), NCBI enterobacteria (52), NCBI Archaea (52), NCBI not enterobacteria-includes every non-eukaryotic genome that is not enterobacteria in Genbank (52), pseudomonas database (62), NCBI enterobacteria high quality (52), NCBI viruses high quality (52), NCBI acinomycetota high quality (52). The CDS of genomes mined were defined using MetaGeneMark (63) and annotated via HMM profile searching against the following domain databases: Pfam (35), NCBI_PGAP (64), dbCAN3 (including CAZY) (65, 66), REBASE Gold Standard proteins (61), DefenseFinder (67), TIGRFAM (68), PADLOC (69), CRISPRCasFinder (70), MyRT (reverse transcriptases) (71), KEGG KOFam (72), and dbAPIS (73).

We decided to redefine CDS for the original contigs described in this study (corresponding to Clade C2 and E 5mYOX pathways) using MetaGeneMark for consistency with updated datasets while conducting genome mining searches. Additional annotations were run searching against the following HMM profile databases: Pfam (35), CRISPRCasFinder (70), dbCAN3 (including CAZy) (65, 66), DefenseFinder (67), KEGG KOFam (72), NCBI PGAP (64), PADLOC (69), REBASE gold standard proteins (61), TIGRFAM (68). Sequence datasets of both the screened protein sequences and updated MetaGeneMark defined CDS translations are provided in the supplementary materials. MetaGeneMark defined CDS translations were used for subsequent bioinformatic analyses (MSA, phylogenetic trees, HMM profile generation described above).

MSAs were generated using either Clustal Omega (74), MAFFT (51), or MUSCLE algorithms (75) (Geneious Prime, Dotmatics, Boston, MA) and phylogenetic trees were generated from the alignments using either Geneious Tree builder via Jukes-Cantor distance model (neighbor-joining or UPGMA tree method) (Geneious Prime, Dotmatics, Boston, MA) or FastTree method (76). The protein structures were predicted using AF2 with three recycles and 5 output models (39). The top ranked model was then used as input for structure homolog searches via Foldseek, searching against the PDB structure database (39, 40, 77, 78).

---

**Fig. S1.**
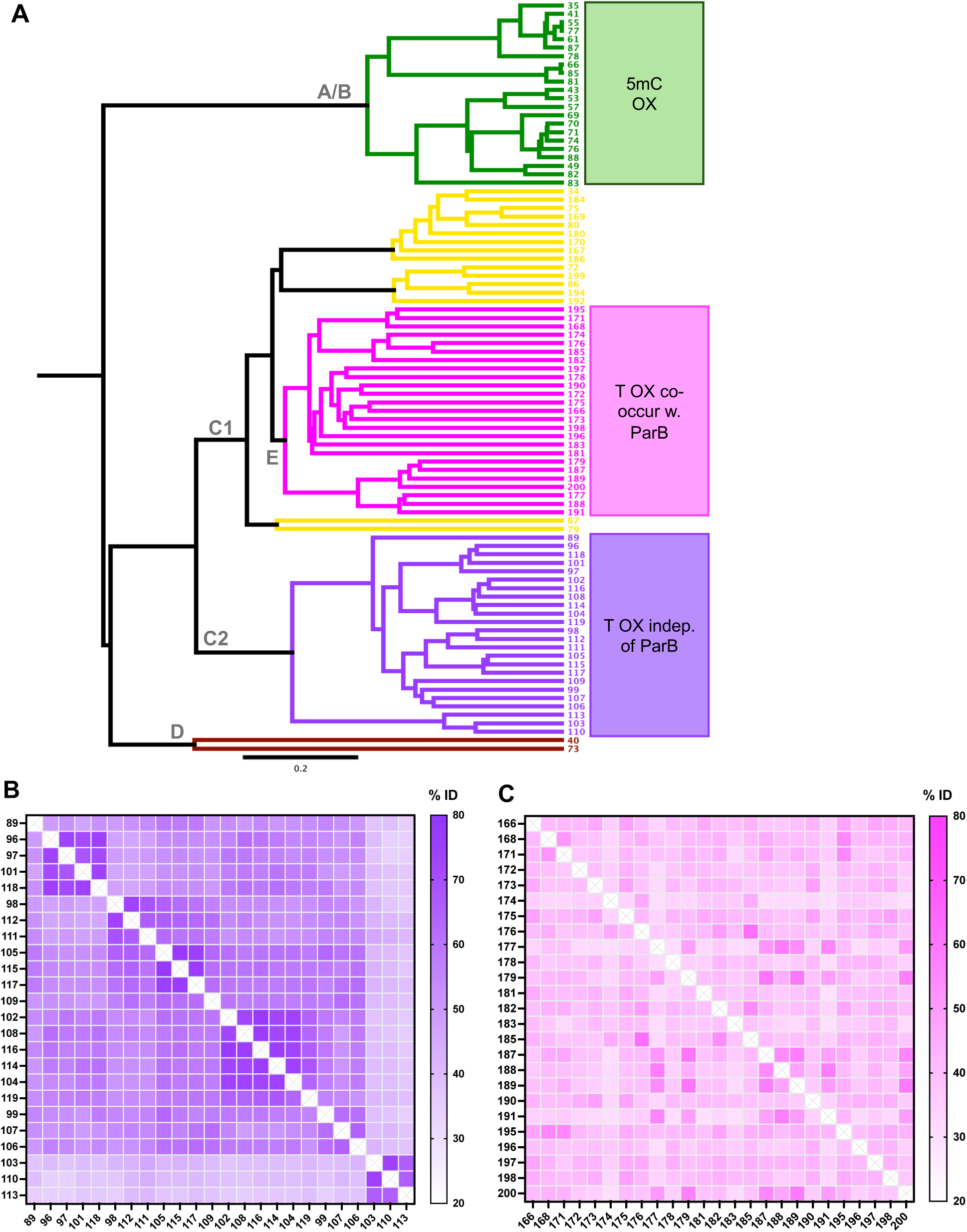
Sequence analysis of 5mYOXs. *(A)* Phylogenetic tree of phage-encoded 5mYOXs from Clades A-E from *Burke et al.* (7) and this study. Tree was generated using Jukes-Cantor distance model and UPGMA tree method from MAFFT-calculated MSAs in Geneious Prime. Heatmaps representing % identity matrix of *(B)* Clade C2 and *(C)* Clade E homologs calculated from MSAs.

**Fig. S2.**
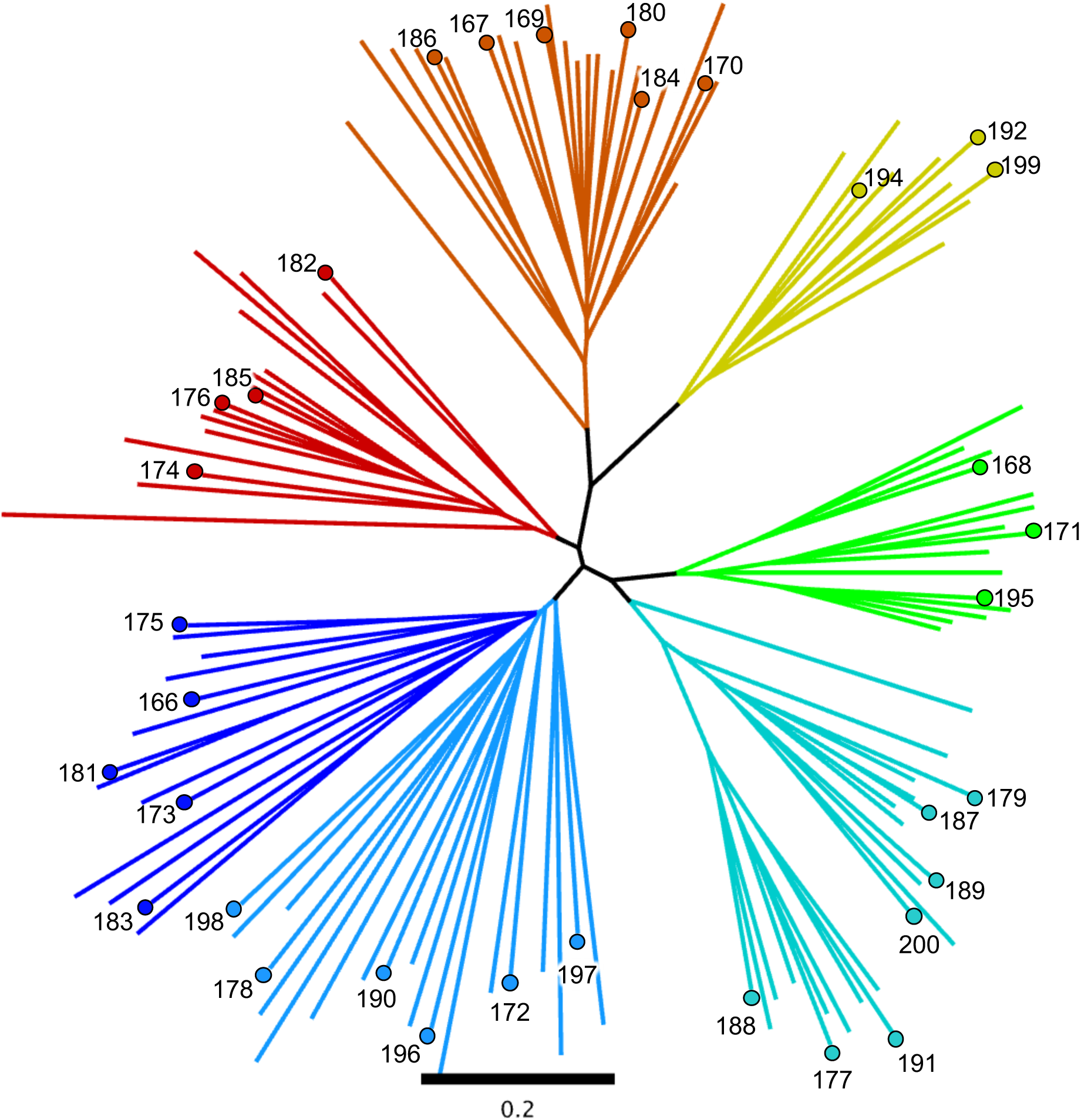
Phylogenetic tree of Clade C1 5mYOXs. Clade C1 homolog prioritization to select diverse representatives co-localizing with ParB that are between 26 and 62 % identical (Materials and Methods). Individual homologs selected are represented with a dot and corresponding number.

**Fig. S3.**
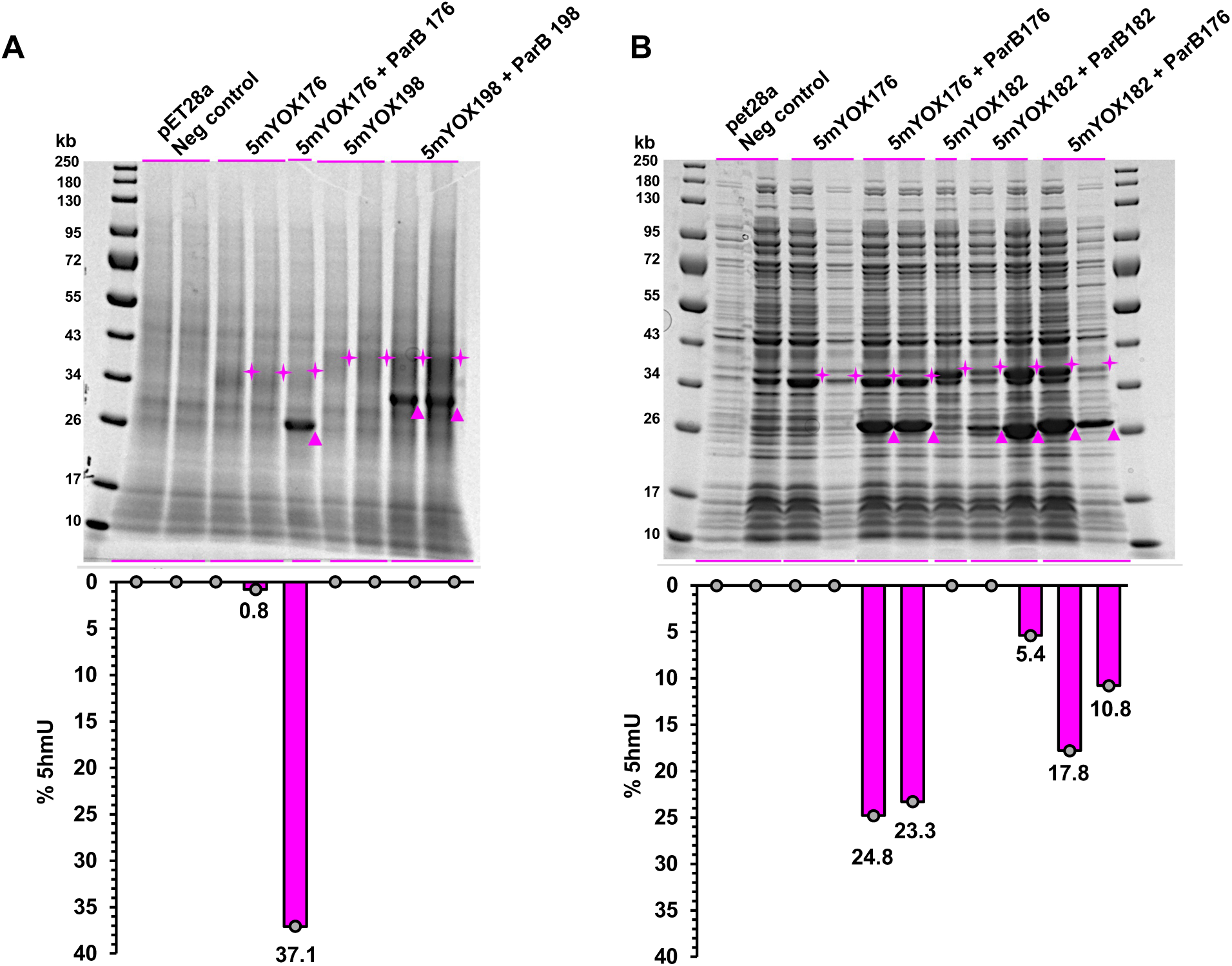
SDS-PAGE analysis of in vivo co-expression of Clade E homologs. Expression conducted in (*A*) 2 mL and (*B*) 0.5 mL cultures. After overnight induction, samples were taken for SDS-PAGE (5mYOX (diamond); ParB (triangle)), and remaining cultures processed for gDNA extraction, digestion, and UHPLC analysis to detect 5hmU. Bar graphs show % 5hmU activity corresponding to expression lanes.

**Fig. S4.**
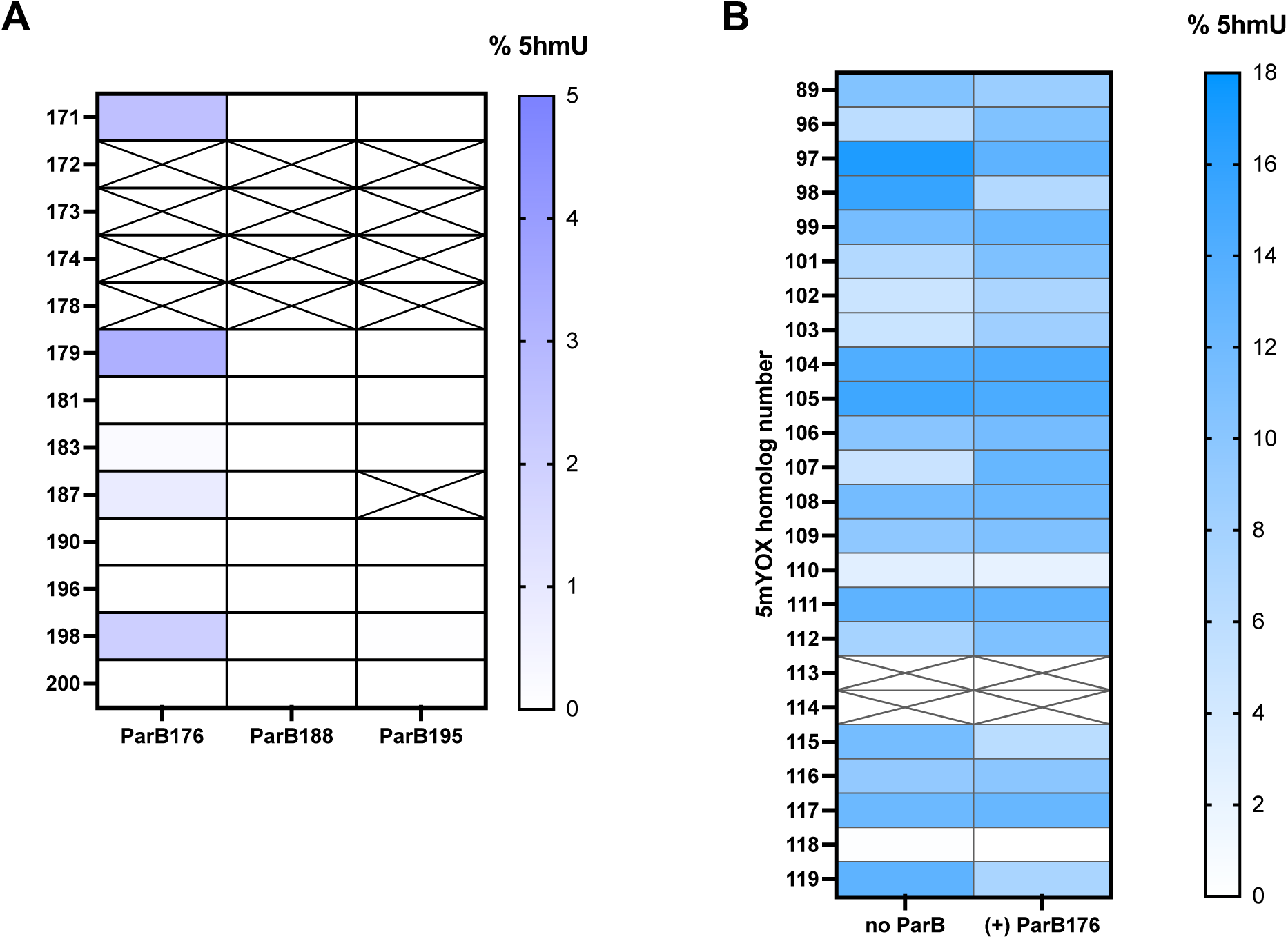
In vivo activity of non-cognate 5mYOX-ParB co-expressions. UHPLC-based activity of (*A)* Clade E 5mYOXs with ParB176, 188, or B195 and *(B)* Clade C2 5mYOXs co-expressed with ParB176. “No ParB” data are extracted from Fig. 4A. Detailed data are in Table S3.

**Fig. S5.**
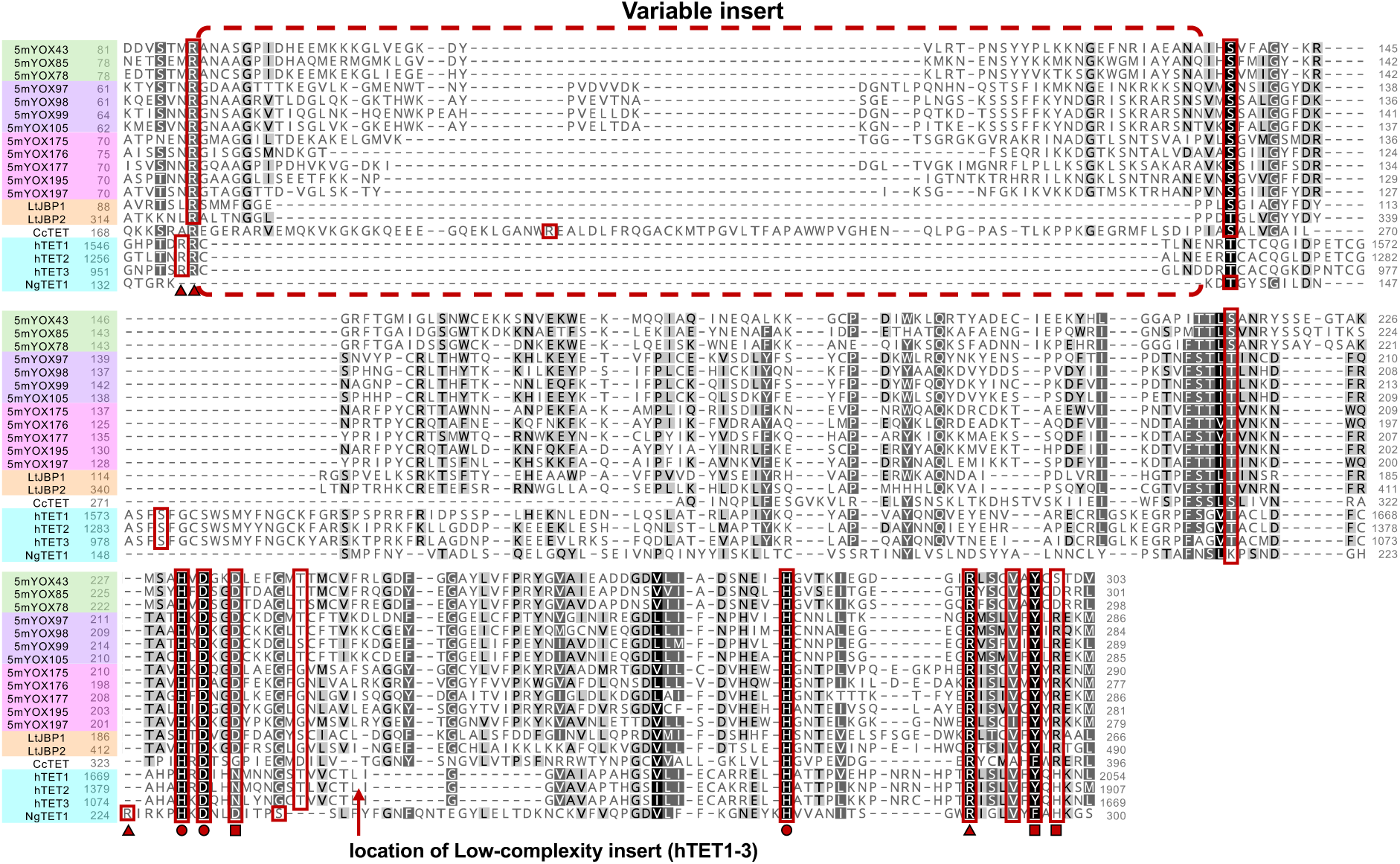
MSA of phage and eukaryotic 5mYOXs. MSA of phage 5mC-dioxygenases (5mYOX 43, 78, 85) (7), Clade C2 T-dioxygenases (5mYOX 97, 98, 99, 105), Clade E T-dioxygenases (5mYOX175, 176, 177, 195, 197), CcTET (UniProt ID: A8P1J0), hTETs 1-3 dioxygenase domain (UniProt IDs: Q8NFU7, Q6N021, and O43151), *Ng*TET1 (UniProt ID: D2W6T1), and *Leishmania tarentolae* JBP1 and 2 (LtJBP1 and 2) dioxygenase domain (UniProt IDs: Q9U6M1 and B6EU02). Accessory domains in hTETs and LtJBPs were removed to enable alignment. Triangle: 2OG interacting residues; circle: Fe(II) coordinating residues; square: nucleobase interacting residues (from crystal structures 4NM6; 4LT5).

**Fig. S6.**
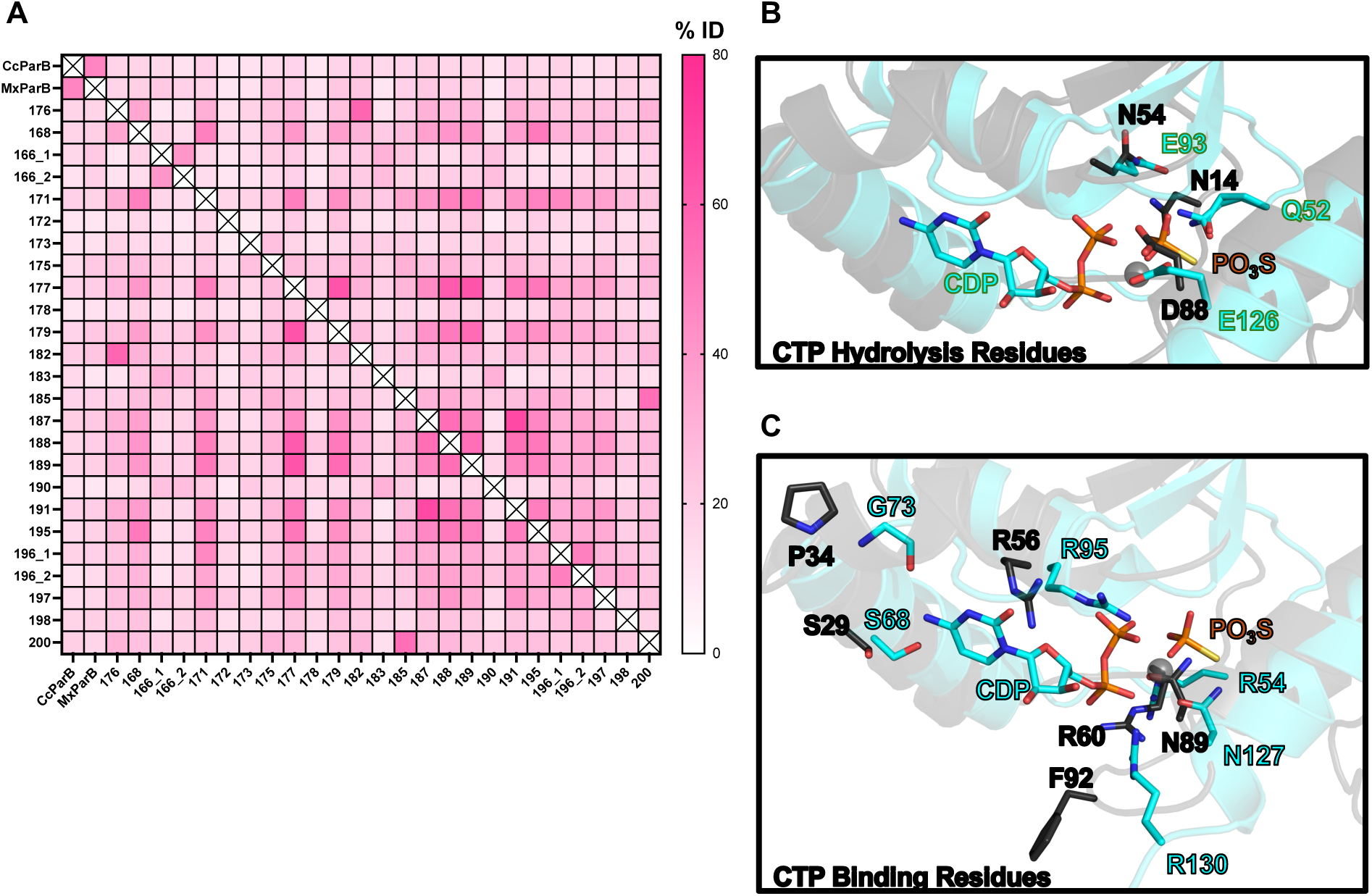
Sequence and structural comparison of 5mYOX-associated ParBs to bacterial MxParB and CcParB. *(A)* Percent identity matrix calculated from MSA of NBDs of 5mYOX-associated ParBs, MxParB, and CcParB (Fig. 6B). *(B & C)* Structural alignment of MxParB (PDB:7BNK, cyan) with AF2-predicted ParB176_2_:5mYOX176_2._ (ParB176 monomer, dark gray). One ParB monomer for each is shown for clarity. CTP (sticks) and Mg^2+^ (sphere) are displayed. Residues implicated in CTP hydrolysis *(B)* or CTP binding *(C)* and the corresponding ParB176 residues (from Fig. 6B) are depicted as sticks, colored by heteroatom.

**Fig. S7.**
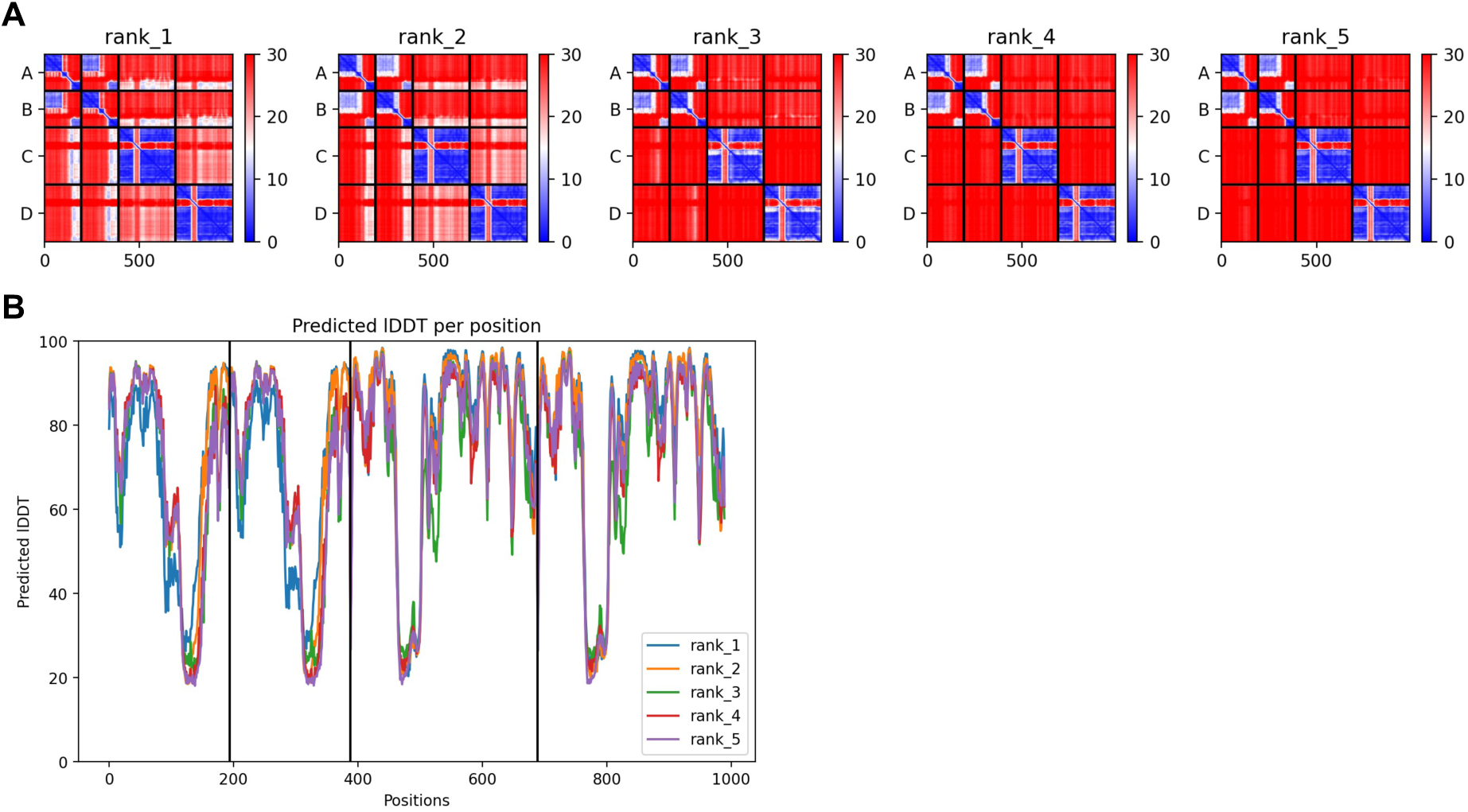
AF2-multimer prediction error plots. *(A)* PAE plots of ParB176_2_:5mYOX176_2_ hetero-tetramer AF2-multimer prediction models, ranked 1-5. Heatmaps show PAE; lower values (blue) = higher confidence between residue pairs. Molecules A/B = ParB176, C/D = 5mYOX176. X-axis shows residue positions for all four chains. Rank 1 model used in Figs. 6A, 6C, 6D. *(B)* <u>P</u>redicted <u>L</u>ocal <u>D</u>istance <u>D</u>ifference <u>T</u>est (pLDDT) plot for the same AF2 predictions. Position numbers on X-axis corresponds to the primary sequence of molecules A-D (ParB176_2_:5mYOX_2_).

**Fig. S8.**
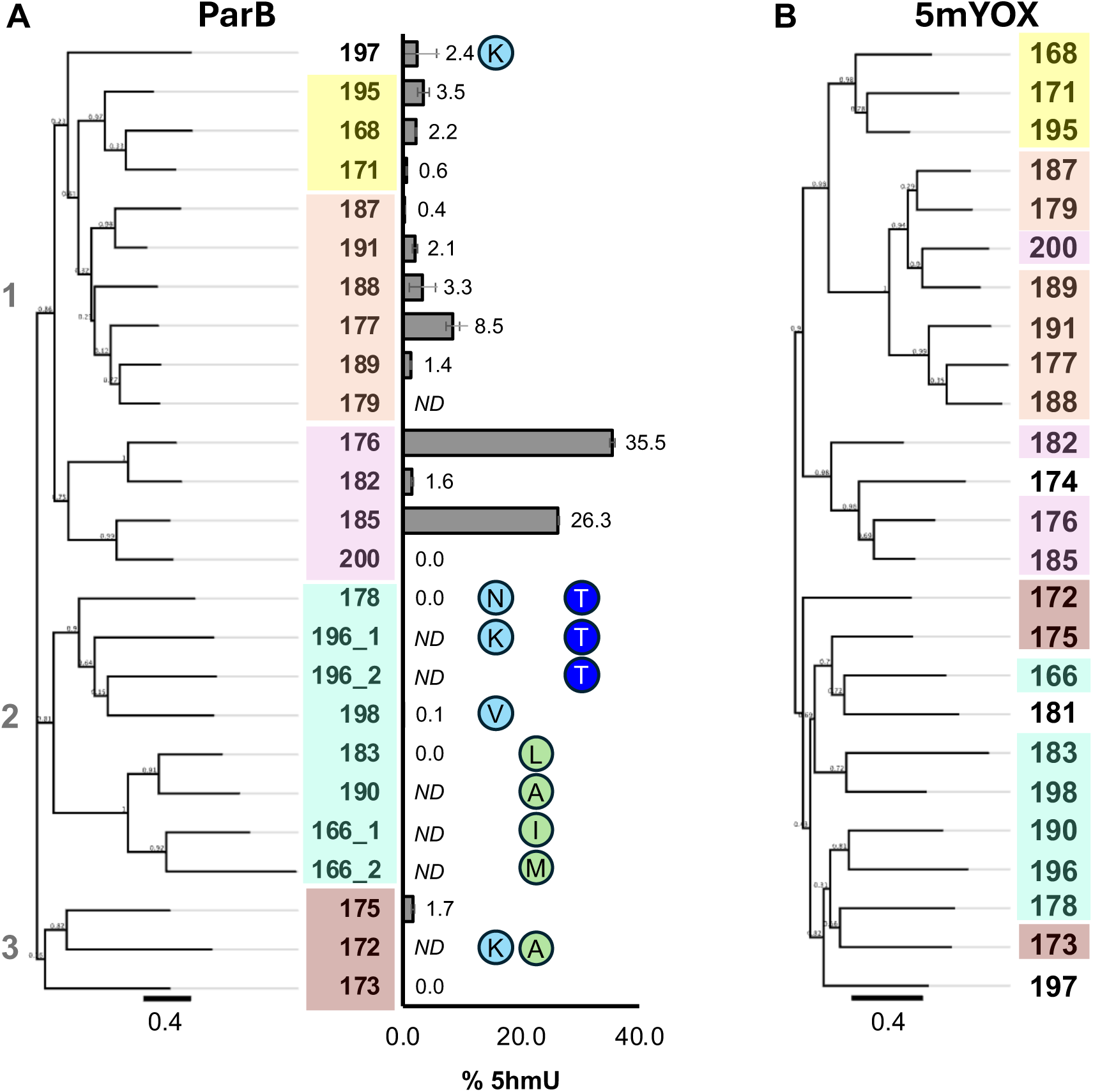
Phylogenetic analysis of Clade E ParB and 5mYOX. Phylogenetic tree of Clade E ParBs *(A)* and 5mYOXs *(B)* were generated using FastTree algorithm from a MAFFT calculated MSA in Geneious Prime. Co-expression activity of 5mYOX-ParB pairs (Fig. 4D and Table S3) is plotted as bars. ND = not determined (unsuccessful assembly). Non-conservation of key ParB active site residue (Fig. 6B) is denoted in colored circles: light blue = R54, green = S68, and dark blue = R95 (MxParB numbering). Contig number colors designate sub-clade organization according to ParB phylogenetic tree.

## Supplemental Table Legends

**Table S1.** Clade C2 in vivo activity data. UHPLC quantification and scalar normalization of nucleoside species from Clade C2 replicate expressions (Fig. 4B).

**Table S2.** Clade E in vivo activity data. UHPLC quantification and scalar normalization of nucleoside species from Clade E replicate expressions (Fig. 4C).

**Table S3.** Non-cognate 5mYOX-ParB in vivo activity data. UHPLC quantification and scalar normalization of nucleoside species from replicate co-expressions of Clade E-ParB (non-cognate) and Clade C2 5mYOX-Clade E ParB (Figs. 4D, S4A, S4B).

**Table S4.** Summary of Foldseek results for phage-encoded 5mYOX enzymes.

**Table S5.** Chimeric 5mYOX in vivo activity data. UHPLC quantification and scalar normalization of nucleoside species from Clade C2 and Clade E chimeric 5mYOXs expressed alone or with ParB (Figs 6C, D).

**Table S6.** Summary of Foldseek results for phage-encoded ParB proteins.

**Table S7.** Genome mining statistics. Summary of genome mining using the following HMM profiles: Clades AB C2 E (5mC- and T-dioxygenases, phage), Clade C2 (T-dioxygenase, phage), and Clade E (T-dioxygenase ParB-dependent, phage) Profiles used for contig searches are listed below each domain label (Fig. 7).

**Table S8.** Scalar normalization values for relative quantification of 5hmU by UHPLC peak areas. SP8 genomic DNA in which every T is replaced with 5hmU was injected on UHPLC in increasing total DNA and monitored at 260 nm. Peak areas were quantified for each injection, and the slope was calculated (increase in peak area divided by increase in sample volume). The scalar value was normalized to dA as described by Pyle *et al*. (23).

